# Sex-dependent effects of intestinal epithelial TLR4 deletion induced before activity-based anorexia

**DOI:** 10.1101/2025.02.05.636626

**Authors:** Colin Salaün, Marion Huré, Charlène Guérin, Christine Bôle-Feysot, Fatima Léon, Sarah Lenoir, Jean-Luc do-Rego, Jean-Claude do-Rego, Ludovic Langlois, David Ribet, Najate Achamrah, Moïse Coëffier

## Abstract

**Rationale:** A role for the microbiota-gut-brain axis in the pathophysiology of anorexia nervosa (AN) has emerged in the last decade. An alteration of intestinal Toll-like receptor type 4 (TLR4) has been reported in the activity-based anorexia (ABA) model with an increase in its expression at the cell surface of colonic epithelial cells. In addition, inducible TLR4 invalidation in intestinal epithelial cells (IECs) was associated with behavioral and energy balance changes in ABA mice. The aim of this study was to assess the intestinal response, e.g. inflammation, gut barrier function and gut microbiota composition, to TLR4 invalidation in IEC in ABA mice.

**Methods:** Male and female Villin-Cre^ERT2^-TLR4 LoxP C57Bl/6 mice were injected with tamoxifen to induce a specific invalidation of TLR4 in IECs (TLR4^IEC-/-^ mice). Then, wild-type (*wt*) and TLR4^IEC-/-^ mice were subjected or not to the ABA protocol which combines an access to a running wheel and a progressively limited access to food. After 12 days, colon samples were collected and the expression of 44 mRNAs encoding proteins involved in inflammatory response, gut barrier function and homeostatic regulation was measured by qPCR. Results were compared by a two-way ANOVA (ABA x TLR4^IEC-/-^). Gut microbiota composition was analysed by 16S rRNA Illumina sequencing.

**Results:** In both male and female ABA TLR4^IEC-/-^ mice, the kinetics of body weight loss was slowed down. In addition, male and female ABA TLR4^IEC-/-^ mice showed an increase and a decrease in food intake, respectively. In males, TLR4 invalidation in IEC was associated with a reduction of *Tlr2*, *Ticam1*, *Myd88*, *Tnfα*, *IκBα*, *Irf3*, *Cxcr3* and *Tgfβ* mRNA expression and fecal calprotectin levels under control conditions but not in response to the ABA model. In females, *Myd88*, *Il6*, *Cxcl1* and *Ccl2* mRNA levels were increased by TLR4^IEC^ invalidation in control mice but not in ABA, except for *Ccl2*. TLR4 invalidation also affected the expression of genes involved in gut barrier function in control and ABA mice in a sex-dependent manner. Male mice exhibited more marked alterations. For instance, male CT TLR4^IEC-/-^ showed a decrease of numerous targets (*Ocln*, *Marveld2*, *F11r*, *Tjp1*, *Cldn7*, *Cldn12*, *Cldn15*). ABA TLR4^IEC-/-^ mice did not exhibit this decrease but other changes were observed such as an increase in *Cldn3* and *Cldn7* mRNA levels. Finally, TLR4^IEC^ invalidation in control mice, but not in ABA, altered the gut microbiota in a sex dependent manner with an increase in the abundance of *Parasutterella* and *Desulfovibrio* genera in females and males, respectively. Interestingly, the ABA model *per se* induced an increase in the abundance of the *Lactobacillus* genus in both sexes, which was not observed in ABA TLR4^IEC-/-^.

**Conclusions:** Our study shows for the first time the impact of inducible TLR4 invalidation in IEC on the intestinal response. We highlighted numerous colonic alterations regarding epithelial permeability, mucosal inflammation and gut microbiota composition, in control and ABA conditions: all were partially reversed in ABA TLR4^IEC-/-^. TLR4 invalidation in IEC also induced changes in energy homeostasis in response to the ABA model both in female and male mice. Further studies are warranted to deeply evaluate the underlying mechanisms.

## Introduction

Anorexia nervosa (AN) is a severe eating disorder characterized by an abnormally low body weight (body mass index < 18.5 kg.m²), an intense fear of gaining weight and a dysmorphophobia according to the Diagnostic and Statistical Manual of Mental Disorders (DSM-5) (Call *et al*., 2013). The prevalence of AN is increasing. Recent data suggest that approximately 1.4% of women and 0.2% of men will develop AN during their lifetime (Galmiche *et al*., 2019). AN also has the highest mortality rate among psychiatric disorders. In a 5-year follow-up study, 227 out of 5,169 women with AN died, resulting in a 4.4% mortality rate (Auger *et al*., 2021). Patients suffering from AN often exhibit compensatory behaviours. Indeed, from 30% to 80% of individuals with AN have inappropriate exercise pattern (Davis *et al*., 1997; Duclos *et al*., 2013). Interestingly, the activity-based anorexia (ABA) model is commonly employed to study AN and shows numerous pathophysiological features observed in patients (Schalla and Stengel, 2019; Scharner and Stengel, 2021). For instance, altered physical activity pattern (Achamrah *et al*., 2017; Duclos *et al*., 2013), altered brain functions and behaviours (Florent *et al*., 2019; Tirelle *et al*., 2022), intestinal hyperpermeability and inflammatory markers (Belmonte *et al*., 2016; Grigioni *et al*., 2022; L’Huillier *et al*., 2019) have been described both in ABA mice and in patients with AN.

Previous studies highlighted the role of the microbiota-gut-brain axis in the regulation of food-related behaviour and in the pathophysiology of eating disorders, particularly in AN (Borgo *et al*., 2017; Fan *et al*., 2023). In ABA mice, a disrupted gut barrier function and increased intestinal inflammation have been reported. Indeed, the Toll-like receptor 4 (TLR4) mRNA expression was reported to be increased in the colonic mucosa of female ABA mice, particularly at the membrane of epithelial cells (Belmonte *et al*., 2016). In addition, both IL-1α and IL-1β were increased in female ABA mice. TLR4 is known for its effects on the regulation of food intake. Notably after infection (Li *et al*., 2021), it mediates the anorexigenic effects of LPS but its role in food behavior in AN remains poorly documented. Surprisingly, TLR4 knock out mice exhibited a high rate of mortality in response to the ABA model (Belmonte *et al*., 2016). In contrast, specific invalidation of TLR4 in intestinal epithelial cells (TLR4^IEC^) only induced beneficial effects on behavior and/or energy homeostasis but in a sex-dependent manner (Tirelle *et al*., 2022). Male ABA TLR4^IEC-/-^ mice exhibited a limited body weight loss during the initial days of undernutrition, although this effect did not reach significance in females. In addition, while wild-type male ABA mice exhibited higher corticosterone levels and increased immobility time in behavioral test compared to controls. This effect disappeared when comparing ABA TLR4^IEC-/-^ to CT TLR4^IEC-/-^ mice (Tirelle *et al*., 2022). Although the impact of TLR4 depletion in IEC on food behaviour and anxiety responses has been studied, its consequences on intestinal responses in ABA mice remains unknown.

In the present study, we thus aimed to get deeper insights into the effects of intestinal epithelial TLR4 invalidation during the ABA model on behaviour and energy homeostasis and to evaluate its impact on intestinal inflammatory response and gut barrier function both in female and male mice.

## Material and method

### Animal experimentation

The present project was approved by the regional ethical committee for animal experimentation (CENOMEXA APAFIS#36881-2022042014271193 v4) and we obtained the authorization to use TLR4 genetically modified rodent models (DUO8167). For all experiments, female and male mice were fed a standard diet A03-3430 (SAFE) and were housed at the platform for animal behavior (“Service Commun d’Analyse Comportementale” or SCAC, HeRaCLeS unit, Rouen university) at 21°C with inversed light/dark cycle.

### Activity-based anorexia model

Mice were subjected to the ABA protocol, or if not, were housed in standard cages (CT). ABA experiments were conducted as previously described (Salaün *et al*., 2024). Briefly, mice were housed in cages equipped with running wheels (Intellibio, Seichamps, France; software Activiwheel). After a period of acclimatation to the cages with free access to food (day 1 to day 5), a progressive limitation of access to food was induced according to the following schedule: 6-5-4-3-3-3-3-3-3-3-3 hours per day from day 6 to day 16, respectively. Food was given at the beginning of the dark phase. During the protocol, body weight, food and water intake were daily monitored at the end of the light phase.

### Invalidation of TLR4 in intestinal epithelial cells

We crossed CreVillin^ERT2^ mice (A kind gift from Dr Sylvie Robine team, Institut Curie, Paris, France) and TLR4_floxed_ mice (Jackson Laboratory, US) to obtain CreVillin^ERT2^ TLR4_floxed_ mice. Then, we invalidated the TLR4 specifically in intestinal epithelial cells in CreVillin^ERT2^ TLR4_floxed_ mice through five intraperitoneal injections of tamoxifen (TMX; T5648, Sigma). Control mice were injected with PBS. We selected mice homozygous for the floxed TLR4 and expressing Cre recombinase. TMX injections were administered through five daily injections of 1 mg dissolved in a solution of 10% ethanol (VWR) and 90% sunflower oil, as previously described (Marjou *et al*., 2004).

The genotype of mice was checked both after birth and post-mortem as previously described (Tirelle *et al*., 2022). As the second LoxP position remained unknown, we performed the *Tlr4* gene sequencing after excision induced by Cre recombinase to localize LoxP sites (Fig. S1A to S1D). For few mice, the Cre recombinase activity was not detected. These mice were thus excluded from the analyses. As a result, two CT males, two ABA males and one ABA female were removed from the study.

To evaluate the effects of TMX itself, we performed TMX or PBS injections into both female and male C57Bl/6 wild-type mice (Janvier Labs, Le Genest-Saint-Isle, France). Food behavior was evaluated by placing mice (female PBS n=6; male PBS n=6; female TMX n=10 and male TMX n=10) in BioDAQ food and drink intake monitor (BioDAQ, Research Diet, Inc., New Brunswick, NJ, US).

At the end of the experiments (day 16), mice were anesthetized intraperitoneally with ketamine (Boehringer Ingelheim, COVETO, Caen, France; 100 mg/kg) and xylazine (Bayer HealthCare, Puteaux, France; 10 mg/kg) diluted in 0.9% NaCl at the end of the resting period. Blood was then collected from the inferior vena cava on heparin-coated Vacutainer tubes. Subsequently, plasma was obtained by centrifugation at 3000 × *g* for 15 min at 4°C. Distal ileum segments were mounted in a Ussing chamber, and colon segments were snap frozen in liquid nitrogen and stored at -80°C until RNA extraction.

### Immunoassays on plasma and fecal samples

Gut peptides regulating metabolism and food intake, GLP1 and PYY were measured by enzyme immunoassays in plasma samples (Phoenix Pharmaceuticals, Karlsruhe, Germany).

At the end of ABA experiments, feces were collected and immediately stored at -80°C until protein extraction. Then, fecal calprotectin, a marker of intestinal inflammation, was quantified by ELISA (S100A8/S100A, R&D System, Mineapolis, US) following manufacturer’s recommendations as previously described (Salameh *et al*., 2020). Each measurement was performed in duplicate.

### Paracellular ileal permeability

Ileal segments were placed in Ussing chambers (Harvard Apparatus, Holliston, MA, US) and the paracellular permeability was evaluated by measuring the flux of FITC-Dextran (4 kDa; excitation: 485 nm; emission: 535 nm) from the mucosal to serosal sides after 3 hours of incubation, as previously described {L’Huillier *et al*., 2019).

### RT-qPCR

Harvested colons were immediately frozen in liquid nitrogen and stored at -80°C until RNA extraction to perform RT-qPCR on inflammation markers and gut barrier genes. Total RNA were extracted from colon using the TRIzol-Chloroform (Invitrogen, Carlsbad, CA, USA; Merck, Darmstadt, Germany) based on the method described by Chomczynski and Sacchi (Chomczynski and Sacchi, 2006). The quality and integrity of the RNAs were assessed via agarose gel electrophoresis. RNA concentrations were quantified using a Nanodrop 2000 spectrophotometer (Thermo Fisher Scientific, Illkirch, France). RNA samples were then treated with DNase (Promega, Charbonnières-les-Bains, France) to remove any contaminating genomic DNA, followed by reverse transcription using M-MLV Reverse Transcriptase (Invitrogen) to generate cDNA as previously described (Salaün *et al*., 2024).

qPCR targeting 44 markers of interest mainly involved in inflammatory responses (*Nod2*, *Tlr2*, *Tlr4*, *Cd14*, *Ticam1*, *Irf3*, *Myd88*, *Nfκb*, *Nfκb-iα*, *Tnfα*, *Il1β*, *Il4, Il6, Il10, Ifng*, *Tgfβ*, *Cxcl1*, *Ccl2* and *Cxcr3*), gut barrier functions (*Ocln*, *Marveld2*, *F11r*, *Igsf5*, *Cgn*, *Tjp1*, *Tjp2*, *Tjp3*, *Mlck*, *Claudins 1*, *2*, *3*, *4*, *5*, *7*, *8*, *11*, *12* and *15*) and eating behaviour regulation (*Gcg*, *Yy* peptide) were then performed using the SYBR Green technology on a QuantStudio 12K Flex real-time PCR system (Life Technologies, Carlsbad, CA, US) at PRIMCACEN platform (HeRacLeS US51, Rouen University). cDNA levels were normalised using the housekeeping genes *Actb*, *Gapdh* and *B2m.* The sequences and melting temperatures of all oligonucleotides used in the study are detailed as reported (Lefebvre *et al*., 2024).

### Gut microbiota 16S rRNA analyses

Cecal contents were immediately frozen in liquid nitrogen and stored at -80°C until DNA extraction for 16S Illumina sequencing, which was subsequently analyzed using the EasyMAP online platform. Specifically, the QIAamp Fast DNA Stool Mini Kit (QIAGEN) was employed for extraction, as previously described (Breton *et al*., 2021). Subsequently, the DNA samples were sent to the University of Minnesota Genomics Center (UMGC) for Illumina MiSeq sequencing of the V5-V6 region of the 16S rRNA gene, resulting in the generation of 2 x 300 bp sequencing products. The obtained data were then analyzed according to the EasyMAP recommendations (Hung *et al*., 2021; http://easymap.cgm.ntu.edu.tw/). Pair-end filtering was conducted using the DADA2 plugin in QIIME-2 to obtain amplicon sequence variants (ASVs) based on the quality-filtered sequences, with a forward trimming range of 6-300 bp and a reverse trimming range of 20-300 bp. Next, the Silva database, which includes the V5-V6 region, was employed for taxonomic analysis. The alpha and beta diversity indexes were calculated in accordance with the recommendations set forth by the EasyMAP pipeline using QIIME-2. Taxonomy differential abundance analyses were conducted using the LefSe LDA pipeline, with Kruskal-Wallis and Wilcoxon test significant thresholds set at p < 0.05 and a threshold on the logarithmic LDA score for discriminative features set at 2. The results were represented graphically as cladograms and bar charts. Kruskal-Wallis pairwise comparisons were employed to analyze alpha diversity indexes and permanova for Jaccard distance.

### Statistical analysis

All statistical analyses and graphs were performed using GraphPad Prism 9 (San Diego, USA). All graphs are presented as mean ± SEM on plots. Significant results were considered when p<0.05. Kinetics of body weight, running wheel and food intake were analyzed by two-way ANOVA (TLR4^IEC-/-^ X time) followed by multiple comparison test. More specifically, for body weight curve, Bonferroni’s was used to compare the differences between the two, CT vs. ABA and *wt* vs. TLR4^IEC-/-^ as used in the same model (Tirelle *et al*., 2022). For running wheel and food intake, Šídák’s post-tests were used to compare the TLR4^IEC-/-^ effect at the same time. Comparison of area under the curve, cumulative food intake, food intake parameters and ileal permeability was performed by unpaired t-test if normality test passed. Grouped analysis of immunoassay and RT-qPCR data was performed by two-way ANOVA (ABA X TLR4^IEC-/-^) followed by Tukey’s post hoc tests. Kruskal-Wallis followed by Dunn’s post-tests were used to evaluate TMX effects. Fecal calprotectin quantification was performed in duplicate and analyzed using a nested t-test.

## Results

### The invalidation of TLR4 in intestinal epithelial cells affects energy homeostasis in response to the activity-based anorexia model in a sex-dependent manner

As previously described (Tirelle *et al*., 2022), both female and male *wt* ABA mice exhibited a body weight loss at day 16 (-7.3% and -15.3%, respectively, Fig. 1A-B). In control conditions, body weight changes were not affected by TLR4 invalidation in IEC both in females (Fig. 1A-B) and males (Fig. 1C-D). In contrast, body weight loss was differentially modified in TLR4^IEC-/-^ ABA mice according to the sex (both, p(TLR4^IEC-/-^)<0.05). Indeed, female TLR4^IEC-/-^ ABA mice showed a lower body weight loss from day 8 to day 12 compared to *wt* ABA mice (up to 5.2% at day 10, Fig. 1A) while male TLR4^IEC-/-^ ABA mice exhibited a lower body weight loss from day 6 to day 13 (up to 4.6% at day 9, p<0.05, Fig. 1C). To better understand these responses, we analyzed both physical activities and energy intake in ABA mice. The physical activity pattern of ABA mice also showed differences related to TLR4 invalidation and sex (Fig. 2). Female TLR4^IEC-/-^ ABA mice presented a lower total and nocturnal physical activity (both p(TLR4^IEC-/-^)<0.05, Fig. 2A-B) compared to *wt* ABA mice, without impact on diurnal activity encompassing food anticipatory activity (Fig. 2C). In contrast, total physical activity was not impaired in male TLR4^IEC-/-^ ABA compared to *wt* ABA mice (Fig. 2D), while nocturnal and diurnal physical activities were, respectively, increased (p(TLR4^IEC-/-^)=0.0128, Fig. 2E) and decreased (p(TLR4^IEC-/-^)=0.0088, Fig.2F). Concerning food intake, again, female and male mice exhibit different behaviors. While female TLR4^IEC-/-^ ABA mice presented a lower food intake compared to female *wt* ABA mice (p(TLR4^IEC-/-^)<0.0001, Fig. 3C), male TLR4^IEC-/-^ ABA mice showed a slight increase in food intake compared to male *wt* ABA mice (p(TLR4^IEC-/-^)=0.0005, Fig. 3D). These differences were also observed in cumulative food intake (Fig. 3C-D; p<0.0001 for females and p<0.05 for males). Of note, food intake was not impacted by TLR4 invalidation in IEC in female CT mice (Fig. 3A), whereas a slight increase was observed in males (p(TLR4^IEC-/-^)<0.0001, Fig. 3B). As food intake was sex-dependently affected by TLR4 invalidation in IEC, we evaluated the intestinal endocrine response by assessing mRNA expression and plasma concentration of GLP1 and PYY peptides (encoded by the *Gcg* and *Pyy* genes, respectively). In females, the ABA model induced an increase in colonic *Gcg* mRNA expression (p(ABA)=0.0012, Fig. 4A), particularly in TLR4^IEC-/-^ ABA mice {Tukey’s post-test, p<0.05). For *Pyy* mRNA levels, an interaction effect was observed without significant differences in post-tests. In contrast to these changes in gene expression, both plasma GLP1 and PYY levels remained unchanged in female *wt* and TLR4^IEC-/-^ ABA mice compared to their respective controls (Fig. 4C). In males, TLR4 invalidation in IEC was associated with a lower *Gcg* mRNA level both in CT or ABA conditions (p(TLR4^IEC-/-^)=0.0064, Fig. 4C). No difference was observed for PYY mRNA expression. ABA model and TLR4 invalidation in IEC, each alone, are associated with a lower plasma level of GLP1 {Tukey’s post-test, p<0.001 and p<0.05, respectively, Fig. 4D) in males, without additive or synergic effect. In contrast, TLR4 invalidation in IEC impaired plasma PYY concentration (p(TLR4^IEC-/-^)=0.0749 and p(interaction)=0.0223, Fig. 4D). Indeed, a trend for an increase in plasma PYY was observed in male CT mice (p=0.0503) that was blunted in ABA conditions.

**Fig. 1.**
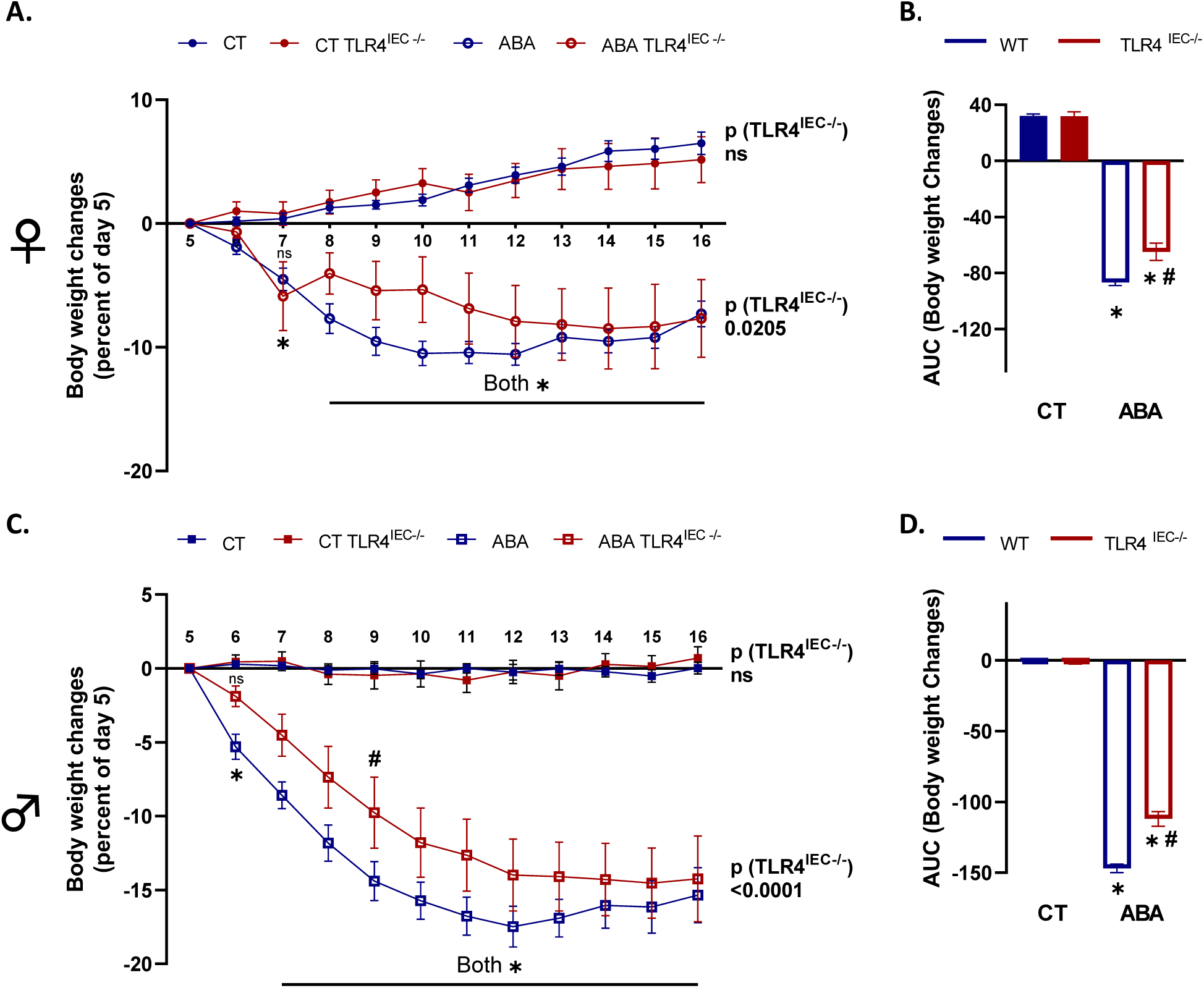
Body weight changes in female and male TLR4^IEC-/-^ mice in response to the ABA model. Body weight changes in female (A, B) and male (C, D) wild-type (*wt*) mice (in blue) and mice invalidated for TLR4 specifically in intestinal epithelial cells (IEC) (TLR4^IEC-/-^, in red) under control conditions (CT, closed symbols) or submitted to the activity-based anorexia (ABA) model (open symbols). (A and C) Data are shown as mean ± SEM and analyzed using two-way ANOVA (TLR4^IEC-/-^ X time). The p value (TLR4^IEC-/-^) is indicated for each condition. Bonferroni’s multiple comparisons test are indicated as *, p<0.05 for ABA vs respective CT and #, p<0.05 for TLR4^IEC-/-^ vs *wt* (n=10-16 per group). (B and D) Area under the curve of body weight changes were analyzed with t-test for females and males. *, p<0.05 for ABA vs respective CT and #, p<0.05 TLR4^IEC-/-^ vs *wt*.

**Fig. 2.**
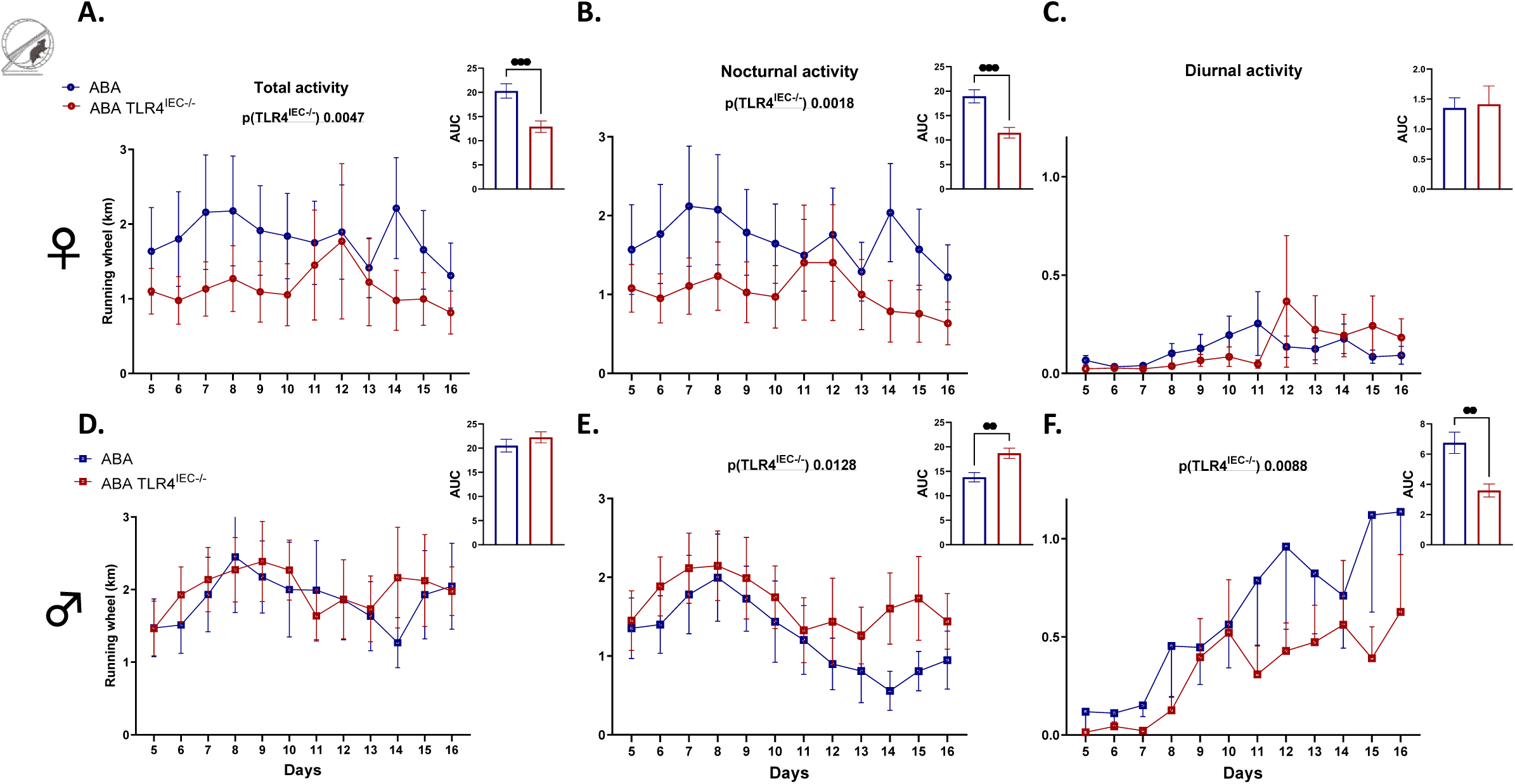
Running wheel activity in female and male TLR4^IEC-/-^ mice in response to the ABA model. Running wheel activities in female (A, B, C) and male (D, E, F) wild-type (*wt*) mice (in blue) and mice invalidated for the TLR4 specifically in the intestinal epithelial cells (TLR4^IEC-/-^, in red) submitted to the activity-based anorexia (ABA) model. Running wheel activity was analyzed as total physical activity (A and D), nocturnal physical activity (B and E) and diurnal physical activity (C and F). Data are shown as mean ± SEM and analyzed using two-way ANOVA (TLR4^IEC-/-^ X time). The significant p value (TLR4^IEC-/-^) is indicated in each graph. Area under the curve is shown in each panel with **, p<0.01 and ***, p<0.001 (Mann-Whitney test, n=10-12 per group).

**Fig. 3.**
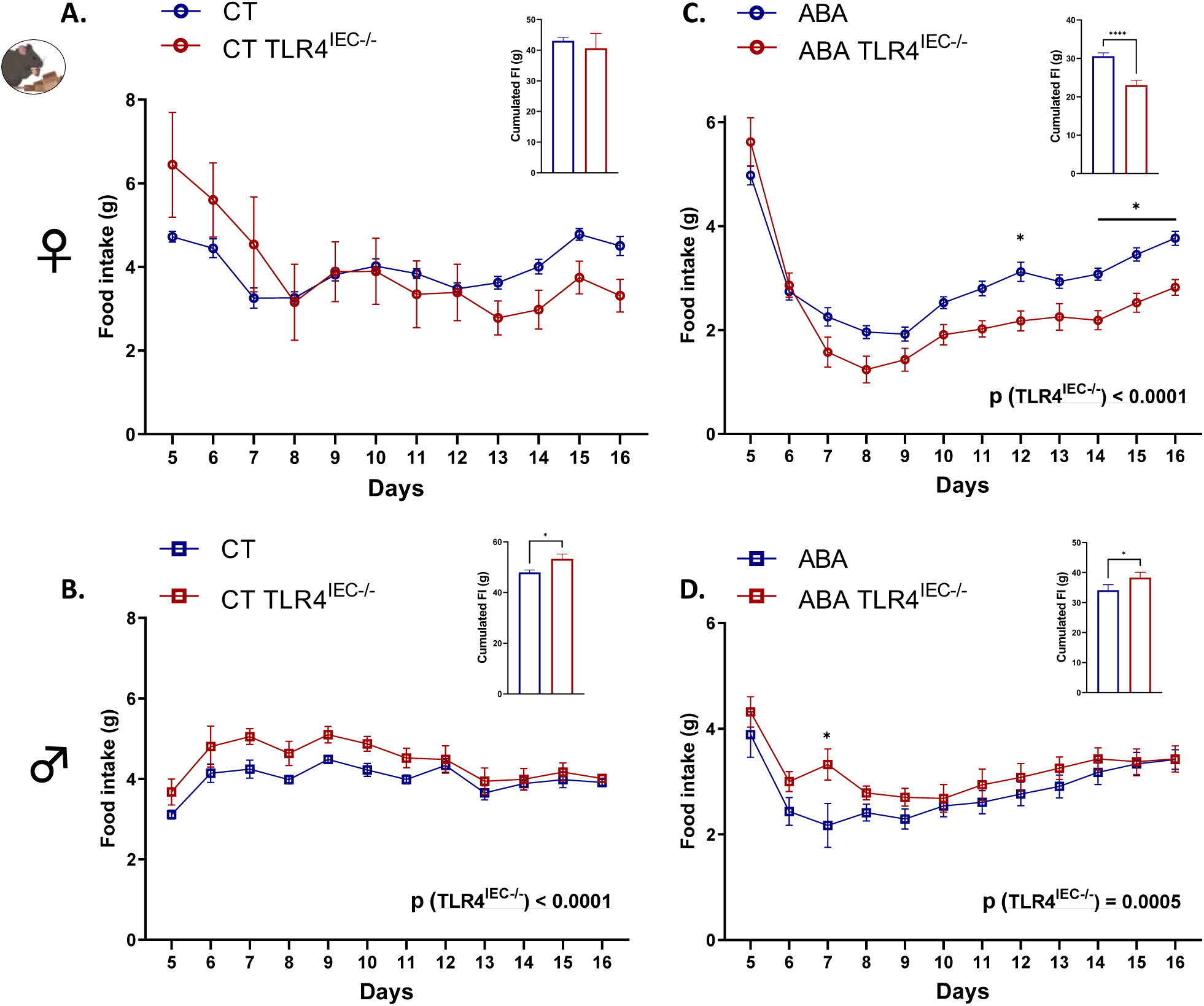
Food intake in female and male TLR4^IEC-/-^ mice in response to the ABA model. Food intake (g) in female (A, C) and male (B, D) wild-type (*wt*) mice (in blue) and mice invalidated for the TLR4 specifically in the intestinal epithelial cells (TLR4^IEC-/-^, in red) submitted to the activity-based anorexia (ABA) model (C, D) or under control conditions (CT; A, B). Data are shown as mean ± SEM and analyzed using two-way ANOVA (TLR4^IEC-/-^ X time). The significant p value (TLR4^IEC-/-^) is indicated in each graph. Šídák’s multiple comparisons test are indicated as *p<0.05 for TLR4^IEC-/-^ vs *wt* (n=10-16 per group). Cumulated food intake is shown in each panel with *p<0.05 and ****p<0.0001 (Mann-Whitney test).

**Fig. 4.**
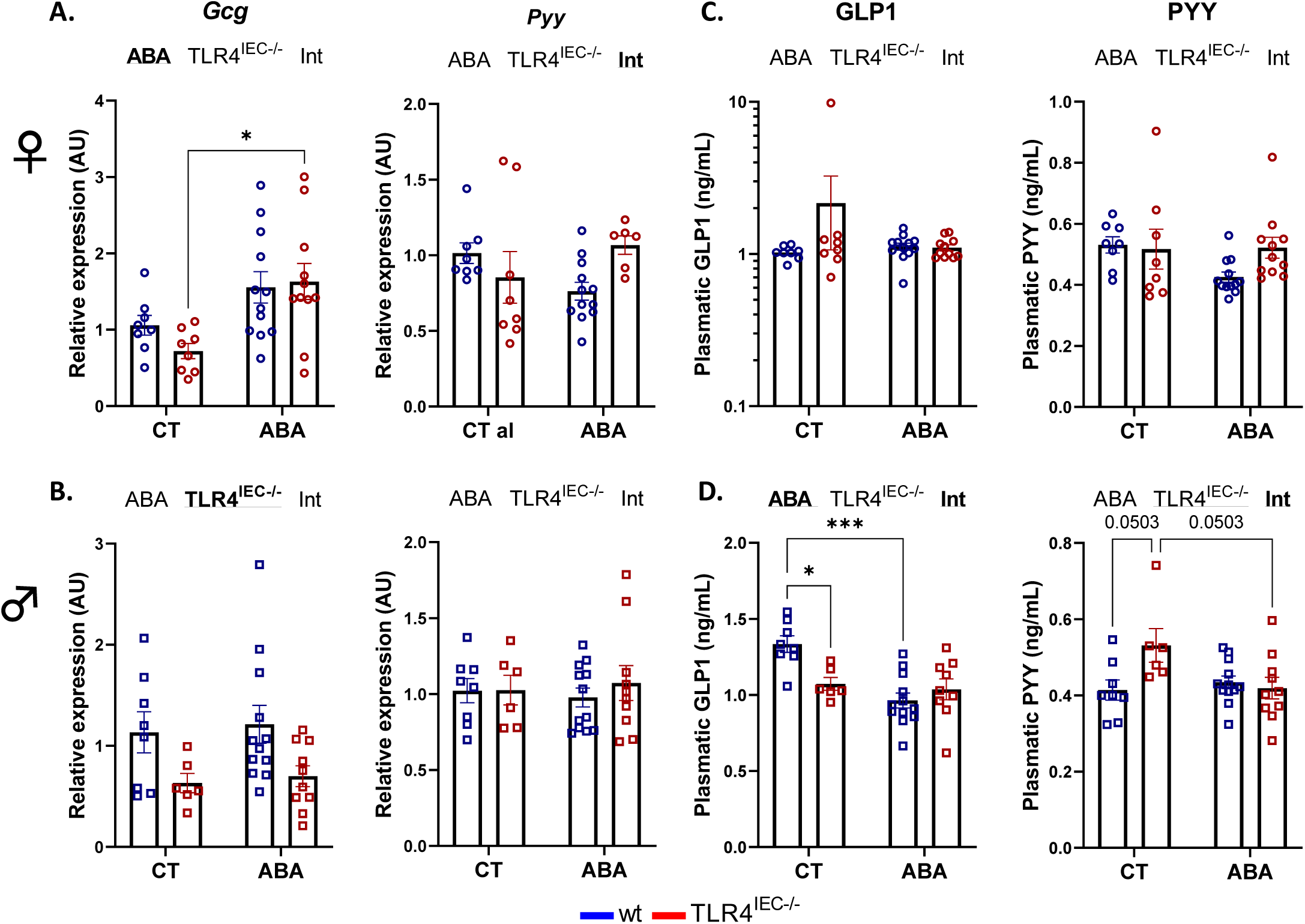
Colonic and plasmatic food intake regulation in female and male TLR4^IEC-/-^ mice in response to the ABA model. cDNA relative levels of *Gcg* and *Pyy* genes in female (A) and male (B) mice. Plasmatic concentration of the anorexigenic hormones GLP1 and PYY in female (C) and male (D) mice. Wild-type (*wt*) mice (in blue) and mice invalidated for the TLR4 specifically in the intestinal epithelial cells (TLR4^IEC-/-^, in red) were submitted to the activity-based anorexia (ABA) model or used under control conditions (CT). Data are shown as mean ± SEM and analyzed using two-way ANOVA (TLR4^IEC-/-^ X ABA). The two-way ANOVA significance (p<0.05 or ABA, TLR4^IEC-/-^ and/or Int for interaction) is indicated by bold and underlined font. Tukey’s multiple comparisons test are indicated as *p<0.05, ***p<0.001 (n=6-12 per group).

As we observed alterations of energy homeostasis in TLR4^IEC-/-^, we investigated the role of TMX injections in control mice (*i.e.* not floxed mice) to discriminate the impact of TMX alone from the impact of TLR4 invalidation, even if TMX injections were performed several days before the beginning of the ABA procedure. Surprisingly, TMX injections limited body weight gain, only in female mice (Fig. S2A). Even if a TMX effect was significant on the kinetics of food intake in both male and female mice (p(TMX)<0.05), cumulative food intake remained unchanged (Fig. S2B-C). Interestingly, eating behavior evaluated in BioDAQ food and drink intake monitor seemed to be affected by TMX alone (Fig. S2C). Particularly, from day 5 to day 16, TMX-injected male mice exhibited higher quantity of pellet waste and higher number of pellets nibbling without consumption (Fig. S2C) than PBS injected mice. We then compared the impact of TMX in *wt* mice to the impact of TLR4 invalidation in IEC under control conditions (*i.e.* not in the ABA model) on entero-hormones expression. In female mice, neither TMX injections nor TLR4 invalidation in IEC modified *Gcg* or *Pyy* mRNA levels, or GLP1 and PYY plasmatic concentrations (Fig. S3). In male mice, only TLR4 invalidation in IEC induced a decrease in plasma GLP1 and a trend for an increase in plasma PYY level (Fig. S3).

### The invalidation of TLR4 in intestinal epithelial cells affects the intestinal barrier function during activity-based anorexia model in a sex-dependent manner

To evaluate intestinal barrier function, we analyzed the expression of 19 gene markers in the colon involved in intestinal permeability (Fig. 5), as well as ileal paracellular permeability (Fig. 6). Again, male mice showed a more marked response compared to female mice. In females, TLR4 ^IEC^ invalidation in CT induced a reduction only in *Cldn1* mRNA expression (Fig. 5A, grey bars) while *Ocln*, *Marveld2*, *F11r*, *Tjp1*, *Cldn7*, *Cldn12* and *Cldn15* mRNA levels were downregulated in males (Fig. 5B). None of these effects were related to TMX injections (Fig. S4A-B). The ABA model *per se* induced slight effects with an upregulation of *Cgn*, *Tjp1* and *Cldn7* mRNA levels and a decrease in *Cldn1* mRNA expression in female mice (blue bars), while only *Tjp1* expression was downregulated in male mice. However, in male TLR4^IEC-/-^ mice, ABA induced an upregulation of *Ocln*, *Marveld2*, *F11r*, *Tjp1*, *Tjp2*, *Tjp3*, *Cldn3*, *Cldn7*, *Cldn12* and *Cldn15* mRNA expression compared to TLR4^IEC-/-^ control mice. In contrast, in female TLR4^IEC-/-^ mice, ABA was associated with an increase in *Cldn2* and *Cldn15* mRNA levels. When comparing TLR4^IEC-/-^ ABA mice to *wt* ABA mice, both in females and males, the mRNA expression of 2 factors was downregulated (*Tjp1* and *Cldn7* in females; *Cgn* and *Cldn5* in males) or upregulated (*Cldn2* and *Cldn15* in females; *Cldn3* and *Cldn7* in males). These data highlight the importance of intestinal TLR4 to modulate gut barrier functions in a sex-dependent manner, even if we did not observe significant differences for ileal paracellular permeability (Fig. 6).

**Fig. 5.**
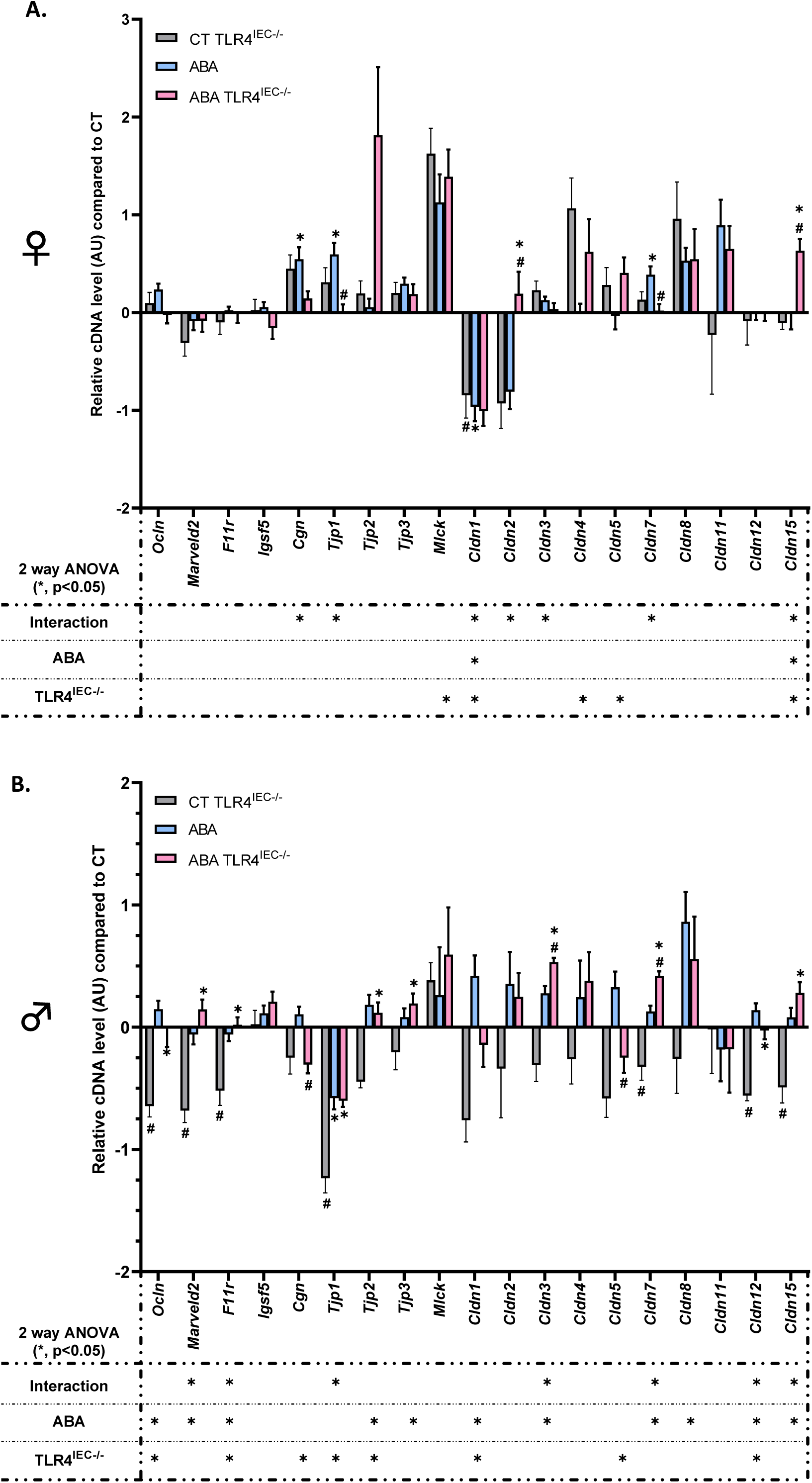
Colonic tight junction protein markers evaluated by RT-qPCR in female and male TLR4^IEC-/-^ mice in response to the ABA model. Relative cDNA levels of tight junction protein markers in control (CT) TLR4^IEC-/-^, activity-based anorexia (ABA) and ABA TLR4^IEC-/-^ groups normalized to wild type (*wt*) CT group in proximal colon of female (A) and male (B) animals. *wt* CT mice or mice submitted to the activity-based anorexia (ABA) model were compared to mice invalidated for the TLR4 specifically in the intestinal epithelial cells (TLR4^IEC-/-^). Data are shown as mean ± SEM bar plot and analyzed using two-way ANOVA (TLR4^IEC-/-^ X ABA). The significance (*p<0.05) were indicated just below the graph. Tukey’s multiple comparisons tests are indicated as * {CT *vs* ABA; CT TLR4^IEC-/-^ *vs* ABA TLR4^IEC-/-^) or # (CT *vs* CT TLR4^IEC-/-^; ABA *vs* ABA TLR4^IEC-/-^). n=5-12 per group except for *Mlck* in males for which CT, n=2.

**Fig. 6.**
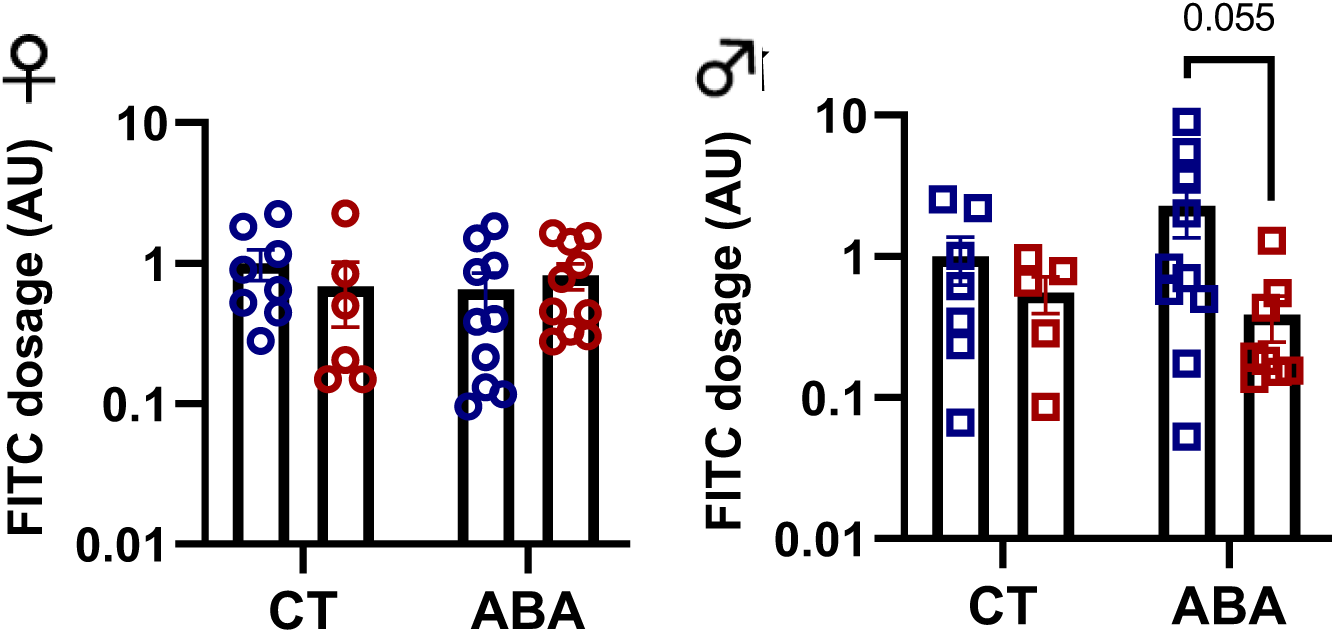
Ileal paracellular permeability evaluated by Ussing chamber in female and male TLR4^IEC-/-^ mice in response to the ABA model. Passage of fluorescent 4 kDa FITC-dextran molecules from luminal to mucosal compartment was evaluated by spectrophotometry on distal ileum segment and express in comparison to the mean of female control (CT). Wild-type (*wt*) mice and mice invalidated for the TLR4 specifically in the intestinal epithelial cells (TLR4^IEC-/-^) were submitted to the activity-based anorexia (ABA) model or used under control conditions (CT). Data are shown as mean ± SEM and analyzed using Mann-Whitney test (n=5-10 per group).

### The invalidation of TLR4 in intestinal epithelial cells affects intestinal inflammatory response during activity-based anorexia model in a sex-dependent manner

To evaluate intestinal inflammatory responses, we measured the expression level of 19 gene markers in the colonic mucosa (Fig. 7). In control and ABA conditions, TLR4 invalidation in IEC decreased TLR4 mRNA level both in female and male mice, as expected (Fig. 7A-B). In control conditions, female TLR4^IEC-/-^ mice only exhibited an increase in *Myd88*, *Il6*, *Cxcl1* and *Ccl2* mRNA expression compared to control wt mice (Fig. 7A) whereas mRNA levels for 8 inflammatory markers were downregulated in male CT TLR4^IEC-/-^ mice (Fig. 7B), including 6 pro-inflammatory (*Tlr2*, *Ticam1*, *Irf3*, *Myd88*, *Tnfα* and *Cxcr3*) and 2 anti-inflammatory markers (*Nfκbiα* and *Tgfβ*). Interestingly, only males showed a lower fecal calprotectin concentration after TLR4 invalidation in IEC (Fig. 7C-D). In addition, TMX injections in control mice did not reproduce these effects (Fig. S5A-B). All of these differences disappeared in ABA conditions (ABA TLR4^IEC-/-^ vs. ABA *wt* mice) both in female and male mice, except for *Ccl2* in female. ABA model *per se* also induced some sex-dependent effects on colonic inflammatory markers. A reduction in *Nod2* and *Tlr4* mRNA expression and an increase in *Irf3* and *Myd88* mRNA levels was observed in female ABA mice. In contrast, male ABA mice showed a decrease in *Nod2*, *Irf3* and *Cxcr3* mRNA levels and an increase in *Ticam1* and *Nf-κb* mRNA expression (Fig. 7A-B). The impact of ABA model in TLR4^IEC-/-^ mice majored the difference according to the sex. Indeed, female TLR4^IEC-/-^ ABA mice only exhibited a reduction in *Nod2* and *Il6* mRNA levels compared to TLR4^IEC-/-^ CT mice (Fig. 7A) that may be explained, respectively, by an ABA or an interaction effect. When comparing male TLR4^IEC-/-^ ABA mice to male TLR4^IEC-/-^ control mice, *Tlr2*, *Cd14*, *Ticam1*, *Irf3*, *Myd88*, *Nfκb, Nfκbia* and *Tgfβ* mRNA levels were increased (Fig. 7B). However, fecal calprotectin was not statistically modified in ABA TLR4^IEC-/-^ mice compared to *wt* ABA mice both in females and males (Fig. 7C-D).

**Fig. 7.**
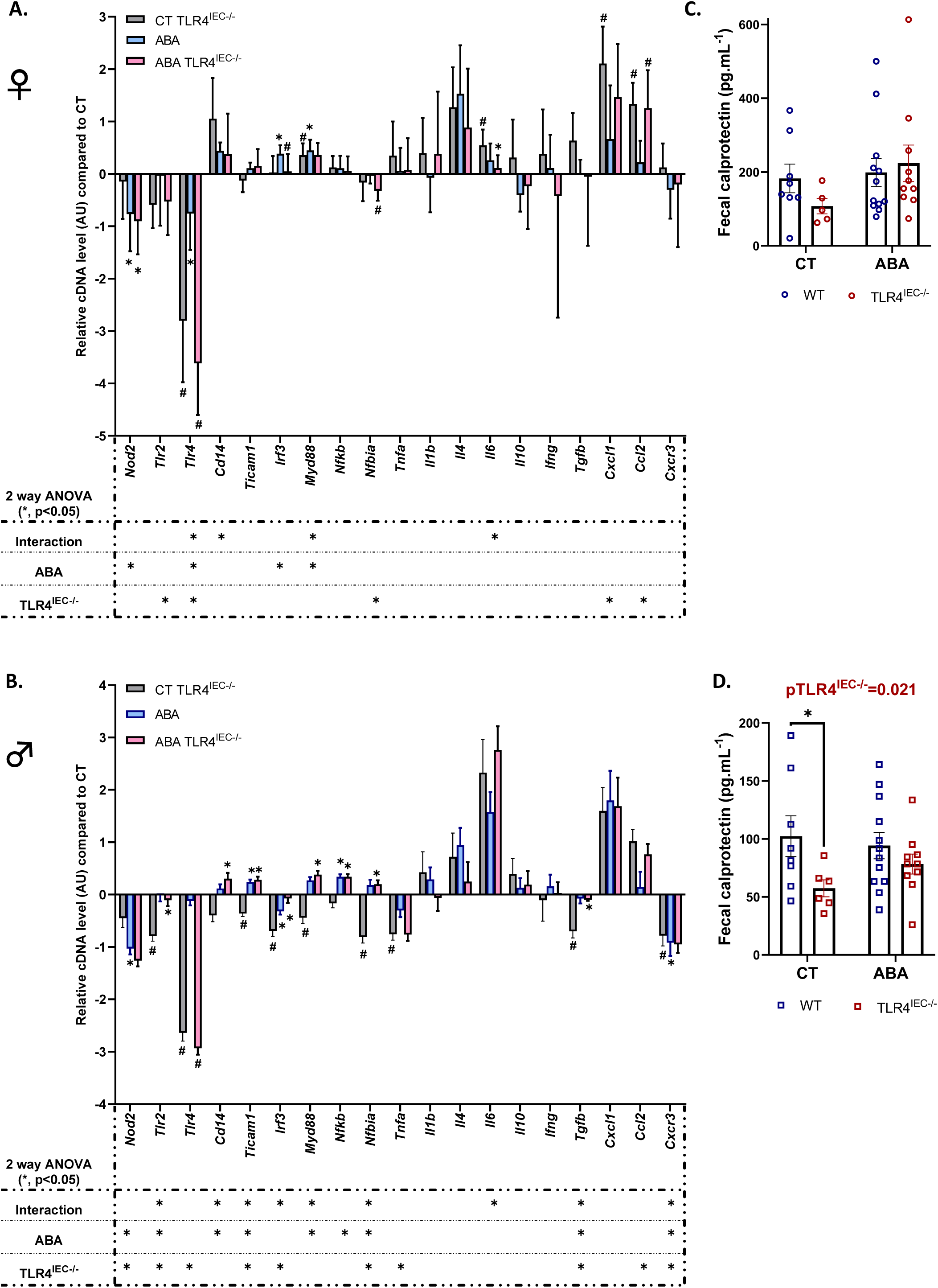
Colonic immunomodulation in female and male TLR4^IEC-/-^ mice in response to the ABA model. Relative cDNA levels of inflammation markers in control (CT) TLR4^IEC-/-^, activity-based anorexia (ABA) and ABA TLR4^IEC-/-^ groups normalized to wild type (*wt*) CT group in proximal colon of female (A) and male (B) animals. *wt* CT mice or mice submitted to the activity-based anorexia (ABA) model were compared to mice invalidated for the TLR4 specifically in the intestinal epithelial cells (TLR4^IEC-/-^). Data are shown as mean ± SEM bar plot and analyzed using two-way ANOVA (TLR4^IEC-/-^ X ABA). The significance (*p<0.05) were indicated just below the graph. Tukey’s multiple comparisons tests are indicated as * {CT *vs* ABA; CT TLR4^IEC-/-^ *vs* ABA TLR4^IEC-/-^) or # (CT *vs* CT TLR4^IEC-/-^; ABA *vs* ABA TLR4^IEC-/-^). Fecal calprotectin in female (C) and male (D). *wt* mice (in blue) and TLR4^IEC-/-^ (in red) submitted to the ABA or not (CT). Two-way ANOVA (TLR4^IEC-/-^ X ABA) and nested t-test *p<0.05, encompassing duplicates values. n=4-12 per group.

### The invalidation of TLR4 in intestinal epithelial cells affects gut microbiota composition during activity-based anorexia model in a sex-dependent manner

We determined microbiota composition by performing V5-V6 16S rRNA gene sequencing on DNA extracted from cecal contents. In female mice, we observed that the total bacterial abundance was reduced in response to TLR4 invalidation in IEC both in CT and ABA mice, whereas the ABA model did not affect it (Fig. 8A). By contrast, in male mice, the bacterial abundance seemed to be reduced by both TLR4^IEC-/-^ and ABA model without additive effects (p(int) < 0.05, Fig 8B), however differences did not reach significance. In male mice, alpha diversity index {Pielou’s evenness) differed between *wt* ABA mice and the CT group, while it remained unaffected in female mice (Fig. S6). The Shannon combined index (richness and repartition) was not significantly affected by TLR4 invalidation and ABA model in both male and female mice (Fig. S6). Concerning the beta diversity (Jaccard distance), the statistical analysis (Permanova) showed differences between the 4 groups in females, CT TLR4^IEC-/-^ vs CT, ABA vs CT, ABA TLR4^IEC-/-^ vs ABA and ABA TLR4^IEC-/-^ vs CT TLR4^IEC-/-^. For males, only ABA TLR4^IEC-/-^ differed from CT TLR4^IEC-/-^ in a significant manner (Fig. S6).

**Fig. 8.**
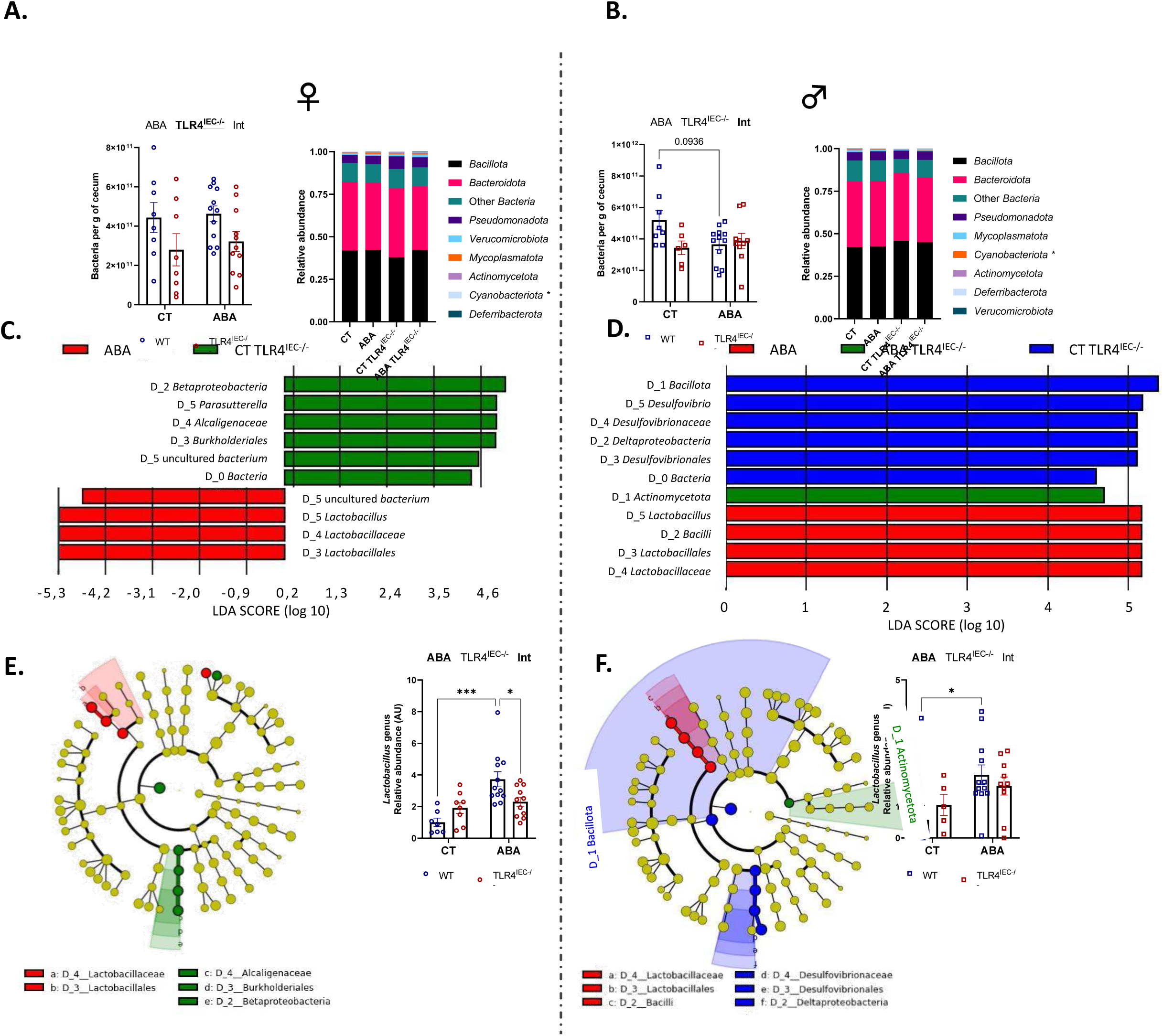
Gut microbiota composition in female and male TLR4^IEC-/-^ mice in response to the ABA model. Gut microbiota composition in cecal content was analyzed in female and male wild type (*wt*) mice and in mice invalidated for the TLR4 specifically in the intestinal epithelial cells (TLR4^IEC-/-^). Mice were under control condition (CT) or submitted to the activity-based anorexia (ABA) model. The total number of eubacteria per gram of cecal content (A and B), the relative abundance of phyla, linear discriminant analysis (LDA) (C and D), cladogram (E and F) and abundance of the *Lactobacillus* genus (E and F) are shown. (C and D), The number preceded by D_ represented the taxa level; domain-0, phylum-1, class-2, order-3, family-4 and genus-5. (A, B, E F). Data were analyzed by two-way ANOVA (TLR4^IEC-/-^ X ABA). Data are expressed as mean ± SEM. Significant (p<0.05) results are indicated in bold and underlined font (ABA, TLR4^IEC-/-^ and Int for interaction). Tukey’s multiple comparison test is indicated as *p<0.05 (n=5-12 per group).

At the phylum level, we observed differences in relative abundances between the groups, that were more pronounced in males (Fig 8A-D). In male CT TLR4^IEC-/-^ mice, the abundance of the *Bacillota* phylum was increased compared to *wt* mice. Similarly, the abundance of *Deltaproteobacteria* class was increased, especially *Desulfovibrionales* order and its associated taxa (*Desulfovibrionaceae* family and *Desulfovibrio* genus; Fig. 8D). Male ABA TLR4^IEC-/-^ mice showed an increase in the abundance of the *Actinomycetota* phylum (Fig. 8B, D). By contrast, in female mice, CT TLR4^IEC-/-^ showed higher levels of *Betaproteobacteria* class (in the *Pseudomonadota* phylum) and *Burkholderiales* order, with higher levels of *Alcaligenaceae* family and *Parasutterella* genus (Fig 8A, C). In response to ABA, both sexes showed elevated levels of the *Lactobacillus* genus (Fig. 8E and F). It is noteworthy that this observed difference in *Lactobacillus* abundance is no longer observed in the ABA TLR4^IEC-/-^ mice when compared to CT TLR4^IEC-/-^ mice, for both sexes (Fig. 8E and F). Of note, the administration of TMX did not affect the total number of bacteria (Fig. S7A). However, male mice that received TMX exhibited a reduction in the total number of observed ASVs and in the Shannon index, in comparison to those that received PBS. The Jaccard distance was significantly different between males injected with TMX or PBS (Fig. S7E, Permanova). Taxa relative abundance (Fig. S7B) revealed some compositional differences, however not aligned with the alterations observed following TMX-induced TLR4^IEC-/-^.

## Discussion

In the present study, we provide evidence that intestinal epithelial TLR4 plays a role in the regulation of energy homoeostasis in response to the activity-based anorexia (ABA) model in a sex-dependent manner. In addition, we show that colonic inflammatory markers, factors involved in the regulation of the gut barrier function and gut microbiota composition are more markedly affected in male mice with intestinal TLR4 deletion compared to female mice in both control and ABA conditions. All these data highlight the putative role of the gut microbiota-host dialogue during anorexia.

Firstly, as we partially previously reported (Tirelle *et al*., 2022), intestinal epithelial TLR4 deletion is associated with an initial limitation of body weight loss in response to ABA model in both sexes. This effect was associated with a decrease in running wheel activity in female mice, despite a decrease in food intake. In contrast, in male mice, TLR4^IEC^ invalidation induced a slight increase of food intake, associated to a reduction of food-anticipatory physical activity. In CT mice, TLR4^IEC^ did not affect body weight, as previously described (Tirelle *et al*., 2022).

The TLR4 inactivation in intestinal epithelial cells induces a disruption of the dialogue between the gut microbiota and intestinal cells, which may interfere with the role of the gut microbiota-gut-brain axis in the pathophysiology of the ABA model (Fan *et al*., 2023). For instance, it has been reported that TLR4 plays a role in mucin production. Cold-inducible RNA-binding protein induced mucin production through the TLR4/NF-κB signaling pathway (Chen *et al*., 2016). Intestinal epithelial TLR4 is essential for the proper development of goblet cells in mice (Grondin *et al*., 2020). In airway epithelial cells, LPS from *Pseudomonas aeruginosa* can induce *Muc5ac* mucin gene expression via TLR4-dependent pathways (Thai *et al*., 2008). Thus, TLR4 signaling contributes to the regulation of multiple mucin genes, including *Muc5ac*, *Muc5b*, and *Muc7*, in response to various inflammatory stimuli and pathogens (Alvestegui *et al*., 2019). Interestingly, female ABA mice exhibited *Muc5ac* alteration in the colonic mucosa (Nobis *et al*., 2018) but also TLR4 expression modification (Belmonte *et al*., 2016). In the present study, TLR4^IEC^ deletion induced a low inflammatory environment in the colon of control male mice that was not observed in female mice. Similarly, factors involved in gut barrier functions were mainly affected in control males but not in females. The novelty of our data is the fact that we used inducible TLR4 ^IEC^ deletion and not a constitutive invalidation as previous studies. For instance, Crame et al reported that constitutive TLR4^IEC^ deletion did not impact gut barrier function and goblet cells both in ileum and colon (Crame *et al*., 2021), but male and female mice were mixed. Similarly, in male mice, non-conditional intestinal TLR4 invalidation did not alter gut barrier function in control conditions, while a gut dysbiosis was observed that might be due to the different origin of mice (*wt* C57BL6/J came from the Shanghai SLAC Laboratory Animal Co. Ltd whereas Villin^+/Cre^/*TLR4^fl/fl^* mice were bred by the authors) (Qi-Xiang *et al*., 2022). *Parasutterellla* and *Burkholderiales* were both increased in TLR4^IEC-/-^ male mice, as previously shown (Qi-Xiang *et al*., 2022), and in female mice (Fig. 8C). In contrast, mice with intestinal epithelial deletion of Myd88, a factor in the TLR4 signaling cascade, showed gut barrier disruption, increased mucus layer and a dysbiosis, particularly a decrease in *Lactobacillus* abundance (Frantz *et al*., 2012) that we did not observe in our model. In our study, we used a model with inducible TLR4^IEC^ invalidation that limits long term adaptations or compensatory mechanisms since TLR4 deletion was induced 22 days before the sampling. Our results could thus reflect an early response to TLR4 deletion. We used TMX to induce TLR4 deletion, which represents a well-established technique, as evidenced by the extensive coverage in the literature (Feil *et al*., 2009). Although some TMX derivatives have been employed, it is important to note that each molecule can activate estrogen receptors (Felker *et al*., 2016). It is of interest to note that a previous study evaluated the efficacy of TMX versus 4-OH TMX showed comparable effects on Cre^ERT2^ activity (Metzger and Chambon, 2001). TMX was initially described to induce no acute toxicity or severe abnormalities in mice (Furr and Jordan, 1984). However, TMX has been documented to delay total transit and alter colonic motility (Bhave *et al*., 2022). Endogenous ERs are expressed not only in gonadal tissues but also in the intestine, kidney, and brain (Barakat *et al*., 2016). These receptors play functional roles in the intestine and are involved in gastrointestinal disorders (Chen *et al*., 2019), which are commonly observed in AN patients, predominantly in females. It is therefore reasonable to ask to what extent the observed sex-dependent results can be attributed to the effects of TMX itself. We have thus evaluated the proper effects of TMX. Even if TMX induced some effects on body weight, eating behaviors and gut microbiota, it did not reproduce the impact of TLR4^IEC^ deletion on inflammatory, gut barrier markers and gut microbiota composition in both males and females.

In mice with TLR4^IEC^ invalidation, our ABA model induced sex-dependent effects on inflammatory markers and gut barrier actors. Again, male TLR4^IEC^ ABA mice showed more pronounced effects than females suggesting a limitation of gut permeability as well as an increase in inflammatory response. To our knowledge, there is no previous study evaluating the impact of inducible TLR4^IEC^ deletion on gut inflammatory responses, barrier function and gut microbiota composition in a pathophysiological condition in both sexes. In a model of acute pancreatitis, constitutive TLR4^IEC^ deletion exacerbated systemic inflammation, ileal permeability and gut dysbiosis in male mice (Qi-Xiang *et al*., 2022), but there was no data in females. In the context of obesity, gut microbiota has been proposed to contribute to the sex-dependent response to high fat diet (Gao *et al*., 2021; Lefebvre *et al*., 2024). It is thus interesting to deeply decipher the role of intestinal epithelial TLR4 deletion in AN-like models and to evaluate its deleterious or beneficial impact. We can speculate that the gut microbiota is differentially affected between male and female by both intestinal TLR4 invalidation and the ABA model mice and that this may contribute to the global sex-specific response to restricted conditions. In addition, the role of *Lactobacillus* that were increased in both ABA model and patients with AN and that were affected by TLR4^IEC^ deletion, especially in females, should be studied further. Another point to discuss is linked to the body weight loss. Indeed, as previously reported (Achamrah *et al*., 2017), male mice lost more weight in response to the ABA model compared to females (approximately, 15% vs 7%). This difference may also contribute to explain sex-dependent response to intestinal epithelial TLR4 deletion on inflammatory responses and gut barrier functions in ABA mice.

In conclusions, our study reinforces the role of the gut microbiota-host interactions in the pathophysiology of AN and highlights the role of intestinal TLR4 in gut homeostasis. Further studies should now decipher the underlying mechanisms involved in the differential response between CT and ABA mice in response to the disruption of the TLR4-mediated microbiota-host dialogue.

## Acknowledgement

We thank Karin Varnier, Sylvie Drouet and Yann Lacoume for their continuous help across the year in the housing of our OGM organisms.

We gratefully thank Sylvie Robine from the Curie Institute, Paris, France for the gift of CreVillin^ERT2^ TLR4_floxed_ mice.

## Figure’s legends

**Fig.S1 – Description of LoxP sites of floxed TLR4 mice**

(A) Genetic structure of *Tlr4* gene and LoxP sites before and after Cre activation. (B) DNA sequence of *Tlr4 mus musculus* gene (S1B) with exons (in capital letters) and introns (in lower-case letters), oligonucleotides used for genotyping (in red) and deleted sequence by Cre (in blue). (C) Analysis of *Tlr4* gene recombination triggered by the Cre recombinase. (D) Evaluation of gut specificity and efficiency of Cre activity on 13 independent animals.

**Fig.S2 – Tamoxifen impact on body weight and food intake from day 5 to 16 in non floxed animals.**

Body weight changes (A) and food intake in g (B) in female (circles) and male (squares). C57Bl/6 wild type mice injected with PBS (in blue) or tamoxifen (TMX, in red). Data are shown as mean ± SEM and analyzed using two-way ANOVA (TMX X time). The significant p value {TMX) is indicated in each graph. Šídák’s multiple comparisons test are indicated as *p<0.05 for TMX vs PBS. Area under the curve (A) and cumulated food intake (B) are shown in each panel with ****p<0.0001 (t-test). Eating behavior (C) was assessed in BioDAQ food and drink intake monitor. Percent of variations of meals quantity, powder produced, time per meal, the number of pellets nibbling without food consumption (meals NaN) and the number of meals were plotted with mean ± SEM and analyzed using two-way ANOVA (Sex X TMX), *p<0.05. Mann-Whitney was used to compare specifically TMX groups compared to their respective PBS group, **p<0.01. n=6-10 per group.

**Fig.S3 – Colonic and plasmatic food intake regulation in female and male is not affected by tamoxifen.**

cDNA relative levels were studied for *Gcg* and *Pyy* genes using the 2^-ΔΔCt^ method and plasmatic concentration of anorexigenic hormones GLP1 and PYY were evaluated by ELISA in female (A) and male (B) mice. C57Bl/6 mice (CT TMX in grey) or CreVillin^ERT2^ TLR4floxed mice (CT TLR4^IEC-/-^ in red) were injected by tamoxifen (TMX), both under *ad libitum* conditions. Data are normalized on their respective controls injected with PBS (CT, dotted line) and shown as bar plots with means ± SEM, and analyzed using Kruskal-Wallis followed by Dunn’s multiple comparisons test, *p<0.05. n=6-10 per group.

**Fig.S4 – Colonic permeability evaluated by RT-qPCR in female and male is not strongly affected by tamoxifen.**

Evaluation of molecular dysfunctions in gut permeability of proximal colon of (A) females and (B) males control mice (CT) to differentiate tamoxifen (TMX) effects alone from invalidation of TLR4 in intestinal epithelial cells (CT TMX and CT TLR4^IEC-/-^ respectively). Kruskal-Wallis followed by Dunn’s multiple comparisons were used here. Significant (p<0.05) test in response to TMX injections, in wild type mice (CT TMX) or mice expressing Cre recombinase (CT TLR4^IEC-/-^) compared to respective control injected with PBS are indicated. Data are expressed as mean ± SEM. n=4-10 /group.

**Fig.S5 – Colonic immunomodulation in female and male is not strongly affected by tamoxifen.**

Evaluation of molecular dysfunctions in immunomodulation of proximal colon of (A) females and (B) males control mice (CT) to differentiate tamoxifen (TMX) effects alone from invalidation of TLR4 in intestinal epithelial cells (CT TMX and CT TLR4^IEC-/-^ respectively). Kruskal-Wallis followed by Dunn’s multiple comparisons were used here. Significant (p<0.05) test in response to TMX injections, in wild type mice (CT TMX) or mice which expressed Cre recombinase (CT TLR4^IEC-/-^) compared to respective control injected with PBS are indicated. Data are expressed as mean ± SEM. n=2-10 /group.

**Fig.S6 – Gut microbiota diversity indexes in female and male TLR4^IEC-/-^ mice in response to the ABA model.**

Alpha diversity (Pielou evenness and Shannon) and Beta diversity (Jaccard distance to CT) were analyzed in female and male wild type (*wt*) mice, and in mice invalidated for the TLR4 specifically in the intestinal epithelial cells (TLR4^IEC-/-^). Mice were under control condition (CT) or submitted to the activity-based anorexia (ABA) model. Data were analyzed using Kruskal Wallis or Permanova pairwise comparisons, respectively.

**Fig.S7 – Gut microbiota composition in female and male in response to tamoxifen (TMX).**

Gut microbiota composition in cecal content was analyzed in female and male wild type mice in response to tamoxifen (TMX) injections. The total number of eubacteria per gram of cecal content (A), the relative abundance of phyla (B), the linear discriminant analysis (LDA) score (C), cladogram (D), alpha diversity (Pielou evenness and Shannon) and Beta diversity (Jaccard distance) (E) are shown. (C), The number preceded by D_ represented the taxa level; domain-0, phylum-1, class-2, order-3, family-4 and genus-5. Data were analyzed using Kruskal Wallis or Permanova pairwise comparisons respectively (E).

