## Supplemental figures for "Sex-dependent effects of intestinal epithelial TLR4 deletion induced before activity-based anorexia"

### Fig. S1 Description of LoxP sites of floxed TLR4 mice

#### (A) Schematic TLR4 gene of Mus musculus

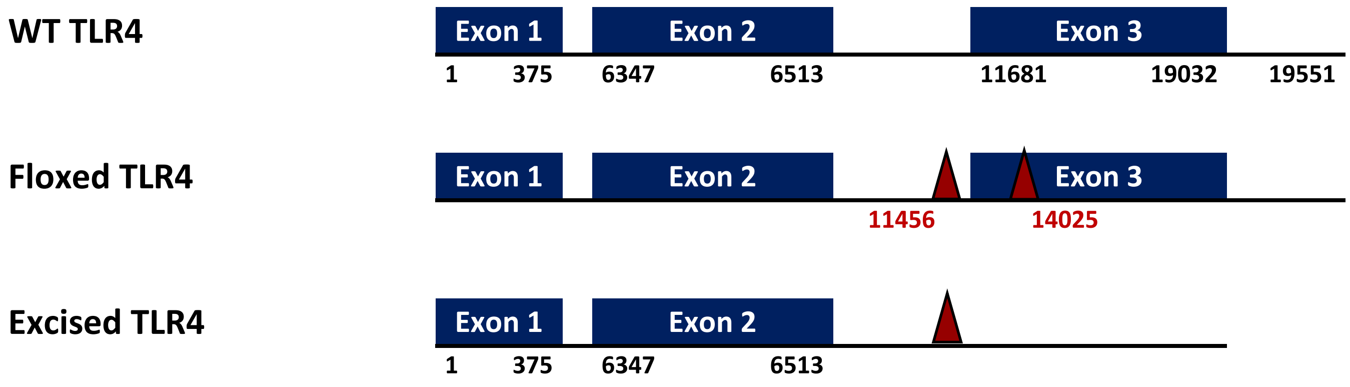


**NCBI sequence identification: NC_000070.7:66745788-66765338 Mus musculus strain C57BL/6J chromosome 4, GRCm39**

#### (B) WT sequence with annotation of primers site and deleted sequence

Exons are in capital. Primer binding sites are in red and deleted sequence in blue.

##### NCBI sequence identification: NC_000070.7:66745788-66765338 Mus musculus strain C57BL/6J chromosome 4, GRCm39

```
AGCTTCCTCTTGCTGTTCCTCCAGTCGGTCAGCAAACGCCTTCTTCCTGTTCTAGTCTTCAGTCTTCTAACTTCCCTCCT
GCGACGGGGCAGATCGATTCTAGAACAAAACCAAAAGTGAGAATGCTAAGGTTGGCACTCTCACTTCCTCTTTGAATATA
GTACTTGCAGAGGGGCACCCACTGGGAGGGAAGAGGCAGGTGTCCCAGGGACTCTGCGCTGCCACCAGTTACAGATCGTC
ATGTTCTCTCATGGCCTCCACTGGTTGCAGAAAATGCCAGGATGATGCCTCCCTGGCTCCTGGCTAGGACTCTGATCATG
GCACTGTTCTTCTCCTGCCTGACACCAGGAAGCTTGAATCCCTGCATAGAGGTAtgtgtcttgatcgcatgtgatcacac
cctttcctgctagcctgccttgtttctcaaaactatccacagctcagagctccctgtgtgtgctctgcttagtttatttt
gcacgaaggagttaaactaaccaaaaacttgagaagccttggcaacaaaaagcctcagtgttaacacagggcaggaacag
gcagccaggggtgtcttgtttcatttaaggcgtctgagtcatgatttagggacttgaaattagtaaaactagtttatagt
cattgttctgtgacatacctgagagtcgttaaagaacttactgaacgtctctgaggccagtattcacgggacgaaagcat
gactgtaatcactgaaaaatgtaagtaggctgtaatttcagggctttctgtgggaactctggccactcagcttttagcgg
tcattccttccctttccaaatcaagtgaaggtagctgtgtcttttctgctgctttcgaagcatctttgagatgctttgag
tggtagctcagcaggtaaggtcagtggctgccaagcctgatgaaaatctgagttcaagcctcaagcctcacaagttagag
gcagggaatctcctcctttaagatgtcttctcacttgcaagtgtctgccttggcaggtgtgtatatgcatgagcacacac
acaaatgaataaagggaacaattgtcttaaatgaaagaatttctattaaaaaataaaacaacaaaacacacaaaaacaca
aagacttttctaagtgattttagtattctgcaactaattctaggagataaagaaatgggaggggtgagggaaggagaggg
acagagcaacttaaaacatcaattagttactgctaaggcagtaactcccgttttggtcgaatactgagtcgtgagtaatc
tgacccatgactcattcttgttttcctcctgcacagaccacgcaattatcttagaagctcacaatagaactgagcaaaca
aggaaggaattcggggtgaggtaggctcagaagctcaaaactggttcaatgagttaagatacatgacattcacatgggga
aaaatactgttaattttaaaaagttataatcacagtatcttgctttctgattcctcagttatgttggcagagatggaatt
tccaatcagtgctacactgagataaaatcccgttgctcttggtgtctggtgtgctttgtcaactctcaaagcttgcttgt
tccttctgtaagccaggtctcagggcccttggccttgtcttcaggagtgattcctgactggtttcctagttcatattcct
ttctatacccacacacagtttcttctttatttgttgttattggtccaggggcttagatttatcaaactactcctttatac
tcttaataactctttggaaccatgatggttgcttcatcctacagggccttagcactgcctaagctaactacacacaccat
catccctcacctaggtcaaggctcaccatgctaaaattatggaatccctgtatatagtttaaaacttcactgttgatcaa
attgaaaaattaagaataaatgcatcaaattagtttcaatgatttttatgcaattaaatatagttatgatgcgtgaaata
taataaaagcatcccacactaacactggctaagcactagcctcaggtctgtctccagccctatggacaggccgaggagaa
catgttctttcctttagccagggtctgtctcacccatgcctgctctgtgtctccagagctctgaaattgctcttttcacc
aggctccataagttaccatggctggctgatgccaagcacgccccacatttccaaattcctgcagctggctggggtgtact
tttttttattagatattttctttatatacatttcaaatgccaccctgaaagttccctataccctccccccaccctgctc
ccctatccacccagtcccacttcttggccctggcgtttccctgtactggagcataaaaagtttgggcctctcttcccagt
gatggctgattaggccatcttctgctacatatgcagctagagatacgagctctgggggtactggttagttcattttggct
ggggtgtactcttgcacaccacactctaccaccatacttttctctggagcccagttgagttgccatgtgaaggaaaacac
aacacacacttggtctacaatcaacaggtaacacaatgttgggtgcagaacctagcatcctaatttttttttattagata
ttttcttaatttacatttcaaatgctatcctcacagcccccctataccctcccctctgccctgctccccaacctacccact
cctgcttcctggctctgccattcccctgtactgtttttgtaaactaatctatgttaaaaatcctccgactcaggagcctc
ttgttcttgtggagacttgaggacccaggataggggaacactaggctgttaaggcaggagtgggtgtgagggtgagggag
caccctcatagaggtagggggggtgggggacggcgagggggtagggggcttgtggagggaaaactgggaagggggataaca
tttgaaatgtaaatgagtaaaataaccaaaaaacaaacaaacaaaatcctcaggtggcagatcttggaggatccaccact
tgaattgacagcctccgactatctgcaatgtgcctctaatgctctcagccatccacaaagagaccttccttactcctgcc
tccctcttcctcttcctcttcccgactcggaagtcccacctactcatctagtgattggtttcctgtaatgtttattaggg
ggaaatcctaccacatagttaagcaattacgaagataccttatgttcaatttttgatacaggaaattagacattcagcaa
catttttgttttactggacattttgatttctcctatgcgtgtttcatatttcatagctatgtgtggcttatagctgcagt
actctaatgtggagctttgatttcaggattatctttttcattttatgtagatttctctgtgaatgtctcctcaggttgat
ttttcttgattgcctcatgtacattttccccttttaccctctccatatgctctttcattgatcatatcattttgtatgttt
gtcttttattttttccaccatttattctcccctttgtgtagaataaacaagaagggagtattactgctgggtttgttagca
tgtcaccaatgcctctcagtggttaacgctaagaccctttagtacagttcctcaggttgtggtgaccttcacccataaaa
ttccttttgttgctacttcttaactataatttttgttatggtgttgaacgataatgtaactatcccctatgcaggatatgt
gatatgtgatcctgtaaatggattgtttgacccttaaatgggtcaaagtccacaggttaagaaccactggcctagatcat
gataggtcttcagttgtatgtgtagtatgtgtgaaaccagtgaaagaatgacttctgaacaccatctgatgtcctcgtgt
tctgcctgtggcttctccatgacagaaggctctgccagtttgtctacatttgttcccacttgttattatttgcttatgtt
cttttctcctttgacatacatatttttttcctttaccacacatttccttgatcagctttccttctgaatctagaatctgt
gtctttgcaactttcgtagttcttattcatgttcttctctgttagctggttctatgagtgcagtgccatcagaaatcatg
taacatgtattcttgtaccacccatggcctttagcagaaaaagcctactatttaacttatacgggctggtgtcccaccaa
ttacacaatatttatcattcattcatccaacaaatgtctattgagcattgagaggtcaccatgtacctttctgagccttg
aagataaatagcaaacaaaaatcatcagagcatcaatgctcatggttcaattgataaatgaaaagcatctggaaaataac
tatataggcaagagatttaccttgtcatcaaaatctgtaaaggaaacaaaagagggtgagagaagaatttctgtctgatg
ccttactctcttagatacattgccttcaaggatccgatgatgagtaccatttagggagatgtgtgtgaagaagcctgttt
```

atgtatgaatcttctgactatatgtgtattaccccacctcttttattttctttgtctttagaggatttttgaagattag
tataaaatacataagttgtaagtaaatgctaatatgtagcaaggaatgaatagtaaccaatgataattaacattaatatt
tatcactttaattaatgcaagctttgagataagctctgatctcatttagccctttgagaattctattgcttttaaataag
agaaaacaaaactcactgggttaagcaaagcattttgccagatgaaatcatataattatgatattacatgaaatgttatg
gtatagggttcacaataaatgtgagaaaacagataaaactagtggagattatgatagagaaaacactcaaccctgagtac
aattttctaccactggaatccatgcactataagacagcctctgatcccaggaccaaactgagaaagtcaatgaatctaag
aacaaaaataattgtcaaaaaataaggcagaatctaggaaatgtctgtatatttttattggtactctccatgtagctgta
tataatgaaaatgatgaattagaacaacaataattttacataaaagtatatacaagcatacattaacatggcttttacat
acaactagcgaggttcacagaagatattataaagtcaaaccagcacacaagcaaaactttgtcccacactcagtattctt
tagttctttgtgtagtgttaaagactcctgcacatgtgtagctgttggccttttacatctcatgtgcaggcagccatgtc
agtgaaactttatgggtgtagcttttgacattaagaatcacagtatcacagtaaagttcgtaacctttggactcataatc
tttcgtcctcctctcagtgatccctgacctgtaggtgttggagttgtattgtaagtgcttccattggcactggactccag
aattctgcattttggttggttgtgatttttttgtcgtgatctctgtttataaagtgggagaaatagtctttcccaagcaa
tagcacagcaattagttaccaaatgccaaatggccaaccctgaaaacatatacataagtaatattatacaaactgaacag
gttctacttatatatgtgggattttatttatacaatatacaatatatatatatcaacaattaatgaagcgggcaacacgg
acttgaaaaacagcaaagacaagggagtaagaaaaaaactttaagagtggaaaaggaaaagtgaagtgatataattataa
tttcaaataatagtaataaaaaagatctactctgtaccaagtggcacacaacacttgttatgaaattaaggttttcagac
ttgagagttatgtaacacctgattctattgtttctcatttaatcataattttgttgtagcagaatgttaacatattgaga
attcaggggatatttttttcttcctgatatgtggaataagatgtcttgcaaatatgaagaggcagataaataaatggagaa
ggatgggtgtgataccatatccccagaatggcaggtattttgggagtccaatgttatctttgactgtatagctaatttaa
ggccagactggtctataggaaagcttgtttcaaccaaaataaatcatgaacgaatgaatgaataggtggacaatatgttg
agtggcatgtacatgtgagagttttatcaccccattattcatctttggagaggagtgggaacacacggttggaaacataa
caattgttgtgtggtatttacaggtaGTTCCTAATATTACCTACCAATGCATGGATCAGAAACTCAGCAAAGTCCCTGAT
GACATTCCTTCTTCAACCAAGAACATAGATCTGAGCTTCAACCCCTTGAAGATCTTAAAAAGCTATAGCTTCTCCAATTT
TTCAGAACTTCAGTGGCTGGATTTATCCAGGTaatgaatgagcttttatgtgatgcagaatgtgaagtagttattttta
tatcattgcattcttggcttagaaaccaaggtggttctaactaaacttccttctgtcatctattcagtagtgctacaac
ttgctgtaaatccttggaaaagctacttttatttaactggtttcagttggatgggccactagataagaatatctaagggc
aattctaacctctacattatttaaaacaatttcattagatatttatgaaccatgtcttatatgttgtatgtctaaactac
agaagaagaatttatagatacaaaacccatactcctaattattaagcaggataaaatcctctttaacaaataagtaagtt
aaagtcttgtccttattattgaacatacagcacaaataaaataaatgttaactaatgctaatactgttgtttataacagt
aagtaataaaatatgtgaaaataagggcaacacactgtgtcctatagaagagtgaatgttttgttatgtgtgtgagagga
tcaggaaagattttgagacatgagtacatatgtaagatacctgaaatattgaaagtagaaaagagagtagagattgaaaa
aaaaactaacttaggagggagatgtaaatgtccaagtaaaacatcaactatgggcaagaaacagttactaagattgtcct
ttctgattcagggcatcttaccatttgttggaacataaaaacttttagccagtatttcaggcgggaagctcaatatattt
tattggttaaaattgctctttgacaatttcatacatctatgtaatgcatacagctactcttaccttcacccacactgagt
tttctctgatcactgttagctctgaccccttccaaaatgtctccaacctatattcataccttcttatttattgtttgacc
cactgattttaaccaggttctctgtgtgaccatagatttagaaaaacctatctgagactagtgaggttaaccatttgata
agcaactaaaaccagtgacggtttctccccaaaaatctaaactttggcagagaagaaatgattccatggtcccctccatg
atcagtaaatatctattggcatgatcagtgcagggaaccacagcttctatgacatcagatttgcaaagtctttgtcatgt
cccacatgtccctcatgtcccacaaatccctcctctctctgtctcttggctcttacatttctatcagattcctcgtcctt
tataatccctgactcttggagagggatttgtgaatgttcattacaggggtgatcacagaactatgttttgcttcttctag
catcttgtacatctaagaatatcctcattcactactgtttactataaagggaagtgacatttgttaaggggtataaatgt
aaatatttagacagaagtctggtactatgctaatttaactaaaccacaataaccaatgccctctctgcacctcaaacatc
agggtcataggcctctctaagcaacatttttttgaacaggttaacagtactagccttggacaaaaatctaatccaagaaag
ctttgttactcctaaaatagttatgccagaatttcagcactggacacatcttgcctggcaggttcatgtaatagttcatc
tgggccatagctggaagagaccagtaatgatttttccccaccagccttcatgacacctttctgctgaaagcaaatcagca
gagagaacattggttgtgcttcagcttcatgtcagtgggttgtactgatcaaggagatccttaggtgttgaagttgaacg
atgaacctcttctctaccatattcctaaagctactggaatgtttcacacatgtgtttttgttctaaaatttagagtatgg
tattaaaagtcttctgcagagcagacaatactgtaaatcattagtgaactagaaaatgtattatactctttacaggagca
tgatagatggagaattccaaaggaagaggaccacagctctgttggtggagcctgtgctttctccaacgtttagcaccatg
tgccctgttgcttgtaacttttcctgagtctctgtcttctctcctagtaaaggaaaatggtaaatctccctccatggtga
aaagttaataaatgagagattattaaaattatttagtgagtttatgagtttgaaaacatgctatcataatcactttatta
aattgtacattctacttatcccagggagatagatttgaagagaactgaggtaagcaggtaaaaaactctaaacagaataa
tctctttttaatatagagaacatagtttttcacccagtataattgagaattgatctaaagtataatgtaagataattcct
taaaggtttggagtttgtattcaggaaaaaggtaagttcctcttcccttagctcacaggatattttgcattagagcaaag
cagacaatctactcctgtgcctttctttaaaaaaaaagataattttcattatgtaatttcaaatgttgtccctttttcctg
gtttccccccctgaaaacccactatcttcacccccctcccccctgctcaccaacacacccacatccacttactggccctggc
attctcttatgttggggcatagaactttcacagcaccaagggcctctcctcccattgatgaccaactaggccattctctg
ttacatatgcagctagagccatgaatcacaccatatgttttctttggttagtggtttagtcccagggagctctgggggta
ctggttagttcatattgttgttcttcctagcactgcaaaccccttcagctccttgggtactttctgtattttattcactg
gggaccctgtgctccgtccaatggatggctgtgagcatccacttctgtatttgtcaggcactggcagaccctctcaggag
acagctatatcaggcttctgtcagaaagctcttgttgatatacacaatagtgcctcaatttgatggttgtttatgggatg
gatccccaggtggcagtctctggatggtcatgccttcagtctcttctccacactttgtctcggtaactcttttcatgggt
attttgttcccacttctaaaaaggattgaagtatgcacactttggccttccttcttcttgagtttcatgtgtttttttgaa
ttgtatcttgggtattctgagcttctgggctaatatccagaattaagtgcatatcatgtgtcttcttttatgactgggtt
acctcactcaggatgatgccctccaggtccattcatttgcctaagaatgtcatagattcactgtttttaatagctgcata
gtactccactgtgcaaatgtaccatatttttttgtatccatttctctgttgagggacatctaggttctttcaagcatctgg
ctattataaataaaactgctatgaacatagtagagcatgtgtccttattacaaggtgaagcatcatctggatatttgcct

tggagtggtattgctggatcctcaggtagtaccatgtccaattttctgaggaaccaccaaactgatttccagagtggtta
tatcagtttacagttctgccagcaatggaagagtgttcctccttctctacatcttgcgagcatctgctgtcacttgagtt
tttgatcttagtcattctgactggtgtgaagtggaatatcagggttgttttgatttgcatttccctgatgactaaggatg
ttaaacatttttttaggtacttttcagtcattcagtattcctcagttgagaattccttctttagttctgtaccccatttt
tcaatatacacaatcataatcatatatgtatgtatatgatttggcaatagaatcctaacagaaagtggaaacttgagaaa
gaatcaaacttagttgcctcatttagaagtggaatgatagaaactcacagaaattaatgggttcccaagatcatgcagga
agaatggagagttaacatggctccatggattcctcttgcgatattctttttaacatacctctacctttgttaaattact
aaggaataaccaaatcacagaccaaaactcttttattacctatgaatactccaaagaaaataggaaaagtgagggaaggt
aattgggttagatttggaagtgactcttttgctaaatgtatctggcatgcatctatgacaacatctgtcatgaatcactg
ttggctgcgtctgagttctgtggctagcttgtctctgtggaagctttacgtagtacagcttacatttatcttggaataaa
atttagaatatttcattgagcttgtgagtctacactattcccactcttgccatacctttatattattcttcctcagtttc
cttgttgcccttcagtcacagagactctgttgtggctcctccgtctggcatgcctgctaactactacaactttggatcg
ctgttttcttcatatattcttcacattcgctcatattgatcattgaaatttccacttacttattctcaagtgtaatctgc
ttttatctggtgagag[illegible]aatattcttaacccattttcttcttcttctataaagcttactc
atgtccctaataatta[illegible]agactcaaaataactagccatcatatatcagtaaagttttgta
Forward primer binding site
aacatttatgccattcttgactcttgacacctatgtgtcattatatatgcctttaaaattaactttcaccagtaatttat
catgactagcaaataa**tgaccacccatattgcctatac**tcattagttgtaaaattatatctatgtctggaaaaaatgcat
aaattaatctaagactactacatatcaactgtctttatgtaccccagttatgatcttgaattgatttttctaatggatt
tgctgcctgacatag**tgtgatagtttatcatcactgtagcaagtgtgaaaatgacaaatctgcagagttcctctcctgct**
**cacaccatcatcacctgttttgctctgtacagttttctctttacaataacatggtatatcatatctgtttgtatcatagt**
**ATGGTAGGGACTGTTA**[illegible]**TTTCAGCAAAAATACATAATTGGTATCTCTTTTGCCCATAGGT**
**GTGAAATTGAAACAAT**[illegible]**CTTACACCACCTCTCAAACTTGATACTGACAGGAAACCCTATC**
Deleted sequence
**CAGAGTTTTTCCCCAGGAAGTTTCTCTGGACTAACAAGTTTAGAGAATCTGGTGGCTGTGGAGACAAAATTGGCCTCTCT**
**AGAAAGCTTCCCTATTGGACAGCTTATAACCTTAAAGAAACTCAATGTGGCTCACAATTTTATACATTCCTGTAAGTTAC**
**CTGCATATTTTTCCAATCTGACGAACCTAGTACATGTGGATCTTTCTTATAACTATATTCAAACTATTACTGTCAACGAC**
**TTACAGTTTCTACGTGAAAATCCACAAGTCAATCTCTCTTTAGACATGTCTTTGAACCCAATTGACTTCATTCAAGACCA**
**AGCCTTTCAGGGAATTAAGCTCCATGAACTGACTCTAAGAGGTAATTTTAATAGCTCAAATATAATGAAAACTTGCCTTC**
**AAAACCTGGCTGGTTTACACGTCCATCGGTTGATCTTGGGAGAATTTAAAGATGAAAGGAATCTGGAAATTTTTGAACCC**
**TCTATCATGGAAGGACTATGTGATGTGACCATTGATGAGTTCAGGTTAACATATACAAATGATTTTTCAGATGATATTGT**
**TAAGTTCCATTGCTTGGCGAATGTTTCTGCAATGTCTCTGGCAGGTGTATCTATAAAATATCTAGAAGATGTTCCTAAAC**
**ATTTCAAATGGCAATCCTTATCAATCATTAGATGTCAACTTAAGCAGTTTCCAACTCTGGATCTACCCTTTCTTAAAAGT**
**TTGACTTTAACTATGAACAAAGGGTCTATCAGTTTTAAAAAAGTGGCCCTACCAAGTCTCAGCTATCTAGATCTTAGTAG**
**AAATGCACTGAGCTTTAGTGGTTGCTGTTCTTATTCTGATTTGGGAACAAACAGCCTGAGACACTTAGACCTCAGCTTCA**
**ATGGTGCCATCATTATGAGTGCCAATTTCATGGGTCTAGAAGAGCTGCAGCACCTGGATTTTCAGCACTCTACTTTAAAA**
**AGGGTCACAGAATTCTCAGCGTTCTTATCCCTTGAAAAGCTACTTTACCTTGACATCTCTTATACTAACACCAAAATTGA**
**CTTCGATGGTATATTTCTTGGCTTGACCAGTCTCAACACATTAAAAATGGCTGGCAATTCTTTCAAAGACAACACCCTTT**
**CAAATGTCTTTGCAAACACAACAAACTTGACATTCCTGGATCTTTCTAAATGTCAATTGGAACAAATATCTTGGGGGGTA**
**TTTGACACCCTCCATAGACTTCAATTATTAAATATGAGTCACAACAATCTATTGTTTTTGGATTCATCCCATTATAACCA**
**GCTGTATTCCCTCAGCACTCTTGATTGCAGTTTCAATCGCATAGAGACATCTAAAGGAATACTGCAACATTTTCCAAAGA**
**GTCTAGCCTTCTTCAATCTTACTAACAATTCTGTTGCTTGTATATGTGAACATCAGAAATTCCTGCAGTGGGTCAAGGAA**
**CAGAAGCAGTTCTTGGTGAATGTTGAACAAATGACATGTGCAACACCTGTAGAGATGAATACCTCCTTAGTGTTGGATTT**
**TAATAATTCTACCTGTTATATGTACAAGACAATCATCAGTGTGTCAGTGGTCAGTGTGATTGTGGTATCCACTGTAGCAT**
**TTCTGATATACCACTTCTATTTTCACCTGATACTTATTGCTGGCTGTAAAAAGTACAGCAGAGGAGAAAGCATCTATGAT**
**GCATTTGTGATCTACTCGAGTCAGAATGAGGACTGGGTGAGAAATGAGCTGGTAAAGAATTTAGAAGAAGGAGTGCCCCG**
**CTTTCACCTCTGCCTTCACTACAGAGACTTTATTCCTGGTGTAGCCATTGCTGCCAACATCATCCAGGAAGGCTTCCACA**
**AGAGCCGGAAGGTTATTGTGGTAGTGTCTAGACACTTTATTCAGAGCCGTTGGTGTATCTTTGAATATGAGATTGCTCAA**
**ACATGGCAGTTTCTGAGCAGCCGCTCTGGCATCATCTTCATTGTCCTTGAGAAGGTTGAGAAGTCCCTGCTGAGGCAGCA**
**GGTGGAATTGTATCGCCTTCTTAGCAGAAACACCTACCTGGAATGGGAGGACAATCCTCTGGGGAGGCACATCTTCTGGA**
**GAAGACTTAAAAATGCCCTATTGGATGGAAAAGCCTCGAATCCTGAGCAAACAGCAGAGGAAGAACAAGAAACGGCAACT**
**TGGACCTGAGGAGAACAAAACTCTGGGGCCTAAACCCAGTCTGTTTGCAATTAATAAATGCTACAGCTCACCTGGGGCTC**
**TGCTATGGACCGAGAGCCCATGGA**ACACATGGCTGCTAAGCTATAGCATGGACCTTACCGGGCAGAAGGAAGTAGCACTG
ACACCTTCCTTTCCAGGGGTATGAATTACCTAACTCGGGAAAAGAAACATAATCCAGAATCTTTACCTTTAATCTGAAGG
AGAAGAGGCTAAGGCCTAGTGAGAACAGAAAGGAGAACCAGTCTTCACTGGGCCTTTTGAATACAAGCCATGTCATGTTC
TGTGTTTCAGTTGCTTTAGAAGAGTATTGATAGTTTCAACTGAACTGAACGGTTTCTTACTTTCCCTTTTTTCTACTGAA
TGCAATATTAAATAGCTCTTTTTGAGAGGTCTTCATTCCAATTTCATCTTCCATTTTATGTCATTTTCTTTTCTTTTTTT
TTTTTATCTAATTCTATAAGAAATATGATTGATACACGCTCACAGATAGCCTGGCCAATCCTAAGAATGCTATATTTATT
AAATACAATTCCTAGTATACTTTTACTTTTATAAATTCAGTTATCGTTTTTCATGCCTTGACTATAAACTAATATCATAA
ATAAGATTGTTACAGGTATGCTAAGAAGGCCCATATTTGACTATAATTTTTTAAGAAAGTATGTAAAATATACTTTGTCA
TATTGTCACTGAATGTCATTCTTAAGTTATTACCTAAGTTATGGATGTCACAGAGTCAGTGTTAAAAATAATTTGGTTGA
TAGAAATATTTTTAATCAGGAGGGAAAAGTGGAGAGGGGTGCAGGAACAGAAATCATGATTTCATCATTTATTCTTGATT
TTTCCGGAAGTTCACATAGCTGAATGACAAGACTACATATGCTGCAACTGATGTTCCTTCTCATCAAGGATACTCTCTGA
AGGACTTGAGAACATTTTGGGGAGGAAGAAAGGTCTAACATCCTTTTCCTTCATCATTCTCATTTCTGGACATGCCTTGT
GAGATGGATGAATGTTGGGAGTACACATTTCTGCTTTCACCTTATTTCAGTCAGCATGAACACTGAATATATAATGTCAT
TTCACAGTGTGTGTGTGTGTGTGTTGTGTATGTACATATATGAACCTGTACATGTGTTTAAGTTTAAAGAGAAAATAGTG
TACAGAGCAGCTCTATATTTGTGATAGGGCTTTAAATAGTTGAGCTAATTCAGAAAAGTATGGAGATTTCTTGGTAAAGG
AAACCAAAGTAGAATCATTACAAGATCTAACAATAAAAATTTTGAAACAATCCTACAAGTAAATATATTGGATTTTCTTG
TCCATTAAGACAATATTCATACTATTGAAATTATGGAAACAACCCTTGGAAGGTTAATGCATAGAGACAGAATGCTATCT
ACTTGCAGTGGAATGTGATTTGACCTTGGAGAAGAAGCAAACCTTGCTACTTGTGAGCAGATGCATAAAGGTGGAGGTTT

TTTATTGTAAGTGAAATATGCCAGGCACAGAAGGAACTGGCCTTTCAGGAACTTTTGATGACATGAGCAAAGTTAGAAAA
AATAATATGCAGAACAATAGAAGAGGAAGACAAA**AGAAAGACAGCCCTAGGATGT**ATTCTTCACAACGATTTTAAACAAT
ATGCTTGAAAGAGAATGAAGTTATTAGTATCAAT[illegible]AATTCCATTCAAACTGGAACATAGC
CACCTAATTATTTGTCTCTTGTTAGCCAAGTGAA[illegible]CATTTTTCTGATATAAAAACCCAAA
TTCTAATGCAGTAAATGTCTTGTCAATCAGCCAG[illegible]GACAGTCTGTGCCCCTTCCCTCTCA
CAGAAACTCCTGTGCACTCTAGCCCACTGCTTCAGGCTACAAGCTAGAAAAGCAAGAAGTGAAAGTGCCACAGTTCTCTA
TGTGGTTAGTGCCAGTCAGGGTCATTCAACTTAAACCATGAGTCATTAAGAAAATACATATGCATGCATGCATTAATGCA
CAGAGTAGTTTATTTATAACAACTCTTTCCATAAAGGGCTGGGGAGTTTTCAACAAAATATAAAGGAACAATTAGTTTAA
TCAAAAGAAAGAAATATAGGCAGAAGAAAGAAATGAAAGAAAGAAAGGAAAGTTTTAACTGTGTATTCCAGGTTTAATTC
TAGAGATCTTCTGGAATTTTAGAGAGTGTGACTTTTGGAGAATTCCTAAACTCATTTTCAGATTATATTACGTATGTGAC
TTGGCCTTCATCTGTCTGAGAGCTAAGAAAGAAATGAAGATCATGCATTTATTATTAGGCCATTACAAACTAATAAATAT
AAAGATAAAAGGGAGACTCTGTGGATGAGTCTCCCTCTTGGCTTTCTTATGGGTAGTCAGAGAGAAGCACTCAGTAGCCT
TATCCTTGACAACATTTTTGTCACATTTGTTTTCCCAGTCTGTAGGACAACAGCAGTCCTTATGACTAAAGTAGATTGTA
TCTTTTTTACCTAGCTTCTATTCATCTGTGTTGTCCTAGCTTCCTTTTTGAGTCTACAGCCTTTGAGAAATCACTAGAAG
TCACTGGAACCTCATGCTTTGACTTGAGGCAGTCCTCATATGTGTTCCTAGGTACTCGAGGGGTCAGTTGGGAGACTGGG
GAGCCATATCTTAACCATCAGCTTTGCTTCCTTGGTGTTGAGCATCATGCCTGACAAAGTAAGCAGACAATGCCTGTATA
CGTGAAGAAGAGGAGAATCATTAATGCATGTTTTCTTGGTGTGCTGTTGTCCTTGATACATTCCAGTTCAGAATCTAAAG
TCCTAGGGATCTTAGCTGTCAACTTAGTTTTCCCTGTCTGTCACTTTGTATGGATGATTTAAATTGCTTCTTCACTTGGT
TGCTTGACACCATGTATTCTAAAATTTTGTGGAAGGTGTGTGTTGGGGGGGGGCGTAGTTCTAACAATAGTGTTCTCTAG
TGGATACATTAAAATCATATTCAGCTAATTAATATTTGATTAAGTTTTGCATGCTATACCGATTTGATAAACATTCACAA
AATCACAGGCTTCAAGATTTTTCTTAACACATCCAAAGTACACAGGCATTAAATGGGCAAAACTAAATATCAAACTGACT
TTATTTAATAGTTTCTCTACTGTTCTCTTTTGTTTTATGTCAAGAGTTGAATGCCACTGTTCTGTATTTTTAATTATTTA
TTGTTTGCTATTGTGAGAATTCAAAGCCAGAACTTTGAGGAGCTGACAGAGGCACTGTGGCCTATGAAGACAGTTTTTGG
AGTTAACAATTTCCTTGGTAACTATGGACTATGTCTCCACACTTCAGCTCTCATATCTGATGGAATAAACTCCTTTCCAG
GAGGCTTCTACTTATGCTAATGCACCCAAGCAAACAAGGAGGCTAATAGAACCAGCTGTTTCTGTCTTTATAGCAATTTC
CCAACATTCTACACTTGAGGATTTCTTCTGTCACATGATTTTTTTCATTGGGCATTCTTTCAATCCTTCATTAAATGGCC
GAGACTTCTCACTAGACCCCAACTCAATGAAATTCTTAAGCTGCTAGCATTGAACAACACTGACTTTTTCAAAGCACCTT
GATAGGGAATTTAAGCTGGACCATCTGAAGCAGGAAAGTCTGTTGTTTTGATGGAATTTCCTAATGGTACCATTGTGGCT
TTATTTTGCCTTGTTAATGTAAGGGATTCAAAGCATTTCAACTTACTACTCATAGTTCAAGCATCTATTTTGCAGATGCA
CTGAAAATTAAGAGATTGGAGAGTTTGTCATATATATTTCCATCATCAACTATTCTAGTTCTTACTAAAGAAGGAGGGTG
CAAAAATTTGAAGGATATGTTAAAGTGCCTTCTATACTTAATGATTCTTCTAGAAAAGGCAAAGTGTTGATCTTGTTCTT
TGTTATGGTATTATATCTTCTCATGGTAATTTGAAAGAAGTTTACATACCAATTTCAGTTTGTTTACCTAGGCCTTGAGA
GTCATTCTACAGTACACGATTAGGCTACTATGAAGACAAAAGAAATCATTGTGGGGAAACTCAGTACAGCTCTAGATTTA
CCTTTTATAATAGATGAATCCCAGAATGATAAAGATCAAGCCTGGCATGATGTTAATTTAGTGGGCTAGGATCCTGGAAA
CCTCCTAAAATAGGACATCCCATGCATTTGGCCTTAGCCAGTGAGGCATCTCTGAGAAAGTGTAGAAAAACTTGCAAGGA
GGTTCAGTGCTCTGAAAGACACAGAGTCAAATGTACATGTAATTCCAGTTCTTCTTTTATATATGTGTACTTTACATAGT
CCCTGAAGTATCGAGAGGCTCAGGTATAGGTGCTACCACCTTGATAGAGTTCACTTAGCCAAAATGCAGAAATGGATGCC
CAGAGAGAATAGATTACTTGTCCTGCATCCTGTAACTTAAAATGTGTTAATAATCATCATAATAAATTCTATCTGCCAAA
TATTTCATATGTGCATGAGACTGTTTTAGTTTAATTATTAAAATTGCTTTCTGATGCAGCTCTTAGCCACATTGTCATTT
CCCATACAATGAAACTGAGACCAAAAAGCAAATTCTCCAATTCCAAGGGTAGAATTCAAGTAATCCTGATATCCAGAGCT
GCTAATTTTTTGCCACACAGTAGACTGCTGCAGTGTCTGGGCTTTTTTGCTGGGGCTCATTCACTCACTAACGGGAGAAT
CCTGTGGACAAGGTCAGCAACTCCCTTACCATCTAGAAATTGAAGGTTTCAAAGGCACTGCATGTGACTTTCCTTGATTT
CTATGGAAATGAAGATGGTCCCTCCTGTGACAGTGCTAAGTGCCGAGTCTGAGTGTAAATGTGCTTTTTGGCACAAATTG
TTCTGTTCTAATAGTGTTGATTATAATTATAAAATAATGTGTTTCTGAAAGGCTGCAAGCAATTCTGGGAATGACAATAA
GGGTTTCGAAACAACATGGTATTTATGTGAGAAGTGTTTTGTTGAAAATTAAACCTGTGTTTAGGAGAAAGgatcctgtt
gtttgctcctaagaaactatcacaccatgtaattaaatcagagccagttggttgccaattggagttcttgtctcacatga
acaatattgtatcacctacaacaaacaagatatgactgaccagaggtagccaagactctttacccaaatcctgtttctct
atcttctcagggcccagaaaaaagatggaaatgcatggtcagtttttttccaaggctgggaattaaccttgtagggtgaa
gccttcctcaagttcatctcagattgtccgtaaggaataggtttttcattcaagggccttttataggaggctgtatctgt
aaataagtgaggaattcaatgtttgagaggctgtcttgacttcctttcttgggaggaaaaacaaaatccttctatgaaga
ttaggaatgtcttcgatgttctcagacctcaaaggcagaaaaaagtatgcagtgtaatttgtttgtatgtatctctctta
aaataatatctaccataacattgtctcccaa

Reverse primer binding site

### (C) Excised site after Cre activity (1768 bp)

Primers binding sites are in red, palindromic sequences of LoxP sites in grey and inserted bases in blue.

**tgaccacccatattgcctatac**tcattagttgtaaaattatatctatgtctggaaaaaatgcataaattaatctaagactact
acatatcaactgtctttatgtaccccagttatgatcttgaattgatttttctaatggatttgctgcctgacatag**atct**ata
acttcgtata**gcatacat**tatacgaagttat**tcgatatcaagcttatcgat**acacatggctgctaagctatagcatggacctt
accgggcagaaggaagtagcactgacaccttcctttccaggggtatgaattacctaactcgggaaaagaaacataatccagaa
tctttacctttaatctgaaggagaagaggctaaggcctagtgagaacagaaaggagaaccagtcttcactgggccttttgaat
acaagccatgtcatgttctgtgtttcagttgctttagaagagtattgatagtttcaactgaactgaacggtttcttactttcc
cttttttctactgaatgcaatattaaatagctcttttgagaggtcttcattccaatttcatcttccattttatgtcattttc
ttttctttttttttttttatctaattctataagaaatatgattgatacacgctcacagatagcctggccaatcctaagaatgct
atatttattaaatacaattcctagtatacttttacttttataaattcagttatcgtttttcatgccttgactataaactaata
tcataaataagattgttacaggtatgctaagaaggcccatatttgactataatttttaagaaagtatgtaaaatatactttg
tcatattgtcactgaatgtcattcttaagttattacctaagttatggatgtcacagagtcagtgttaaaaataatttggttga
tagaaatatttttaatcaggagggaaaagtggagaggggtgcaggaacagaaatcatgatttcatcatttattcttgattttt
ccggaagttcacatagctgaatgacaagactacatatgctgcaactgatgttccttctcatcaaggatactctctgaaggact
tgagaacattttggggaggaagaaaggtctaacatccttttccttcatcattctcatttctggacatgccttgtgagatggat
gaatgttgggagtacacatttctgctttcaccttatttcagtcagcatgaacactgaatatataatgtcatttcacagtgtgt
gtgtgtgtgttgtgtatgtacatatatgaacctgtacatgtgtttaagtttaaagagaaaatagtgtacagagcagctcta
tatttgtgatagggctttaaatagttgagctaattcagaaaagtatggagatttcttggtaaaggaaaccaaagtagaatcat
tacaagatctaacaataaaaattttgaaacaatcctacaagtaaatatattggattttcttgtccattaagacaatattcata
ctattgaaattatggaaacaacccttggaaggttaatgcatagagacagaatgctatctacttgcagtggaatgtgatttgac
cttggagaagaagcaaaccttgctacttgtgagcagatgcataaaggtggaggtttttttattgtaagtgaaatatgccaggca
cagaaggaactggcctttcaggaacttttgatgacatgagcaaagttagaaaaaataatatgcagaacaatagaagaggaaga
caaa**agaaagacagccctaggatgt**

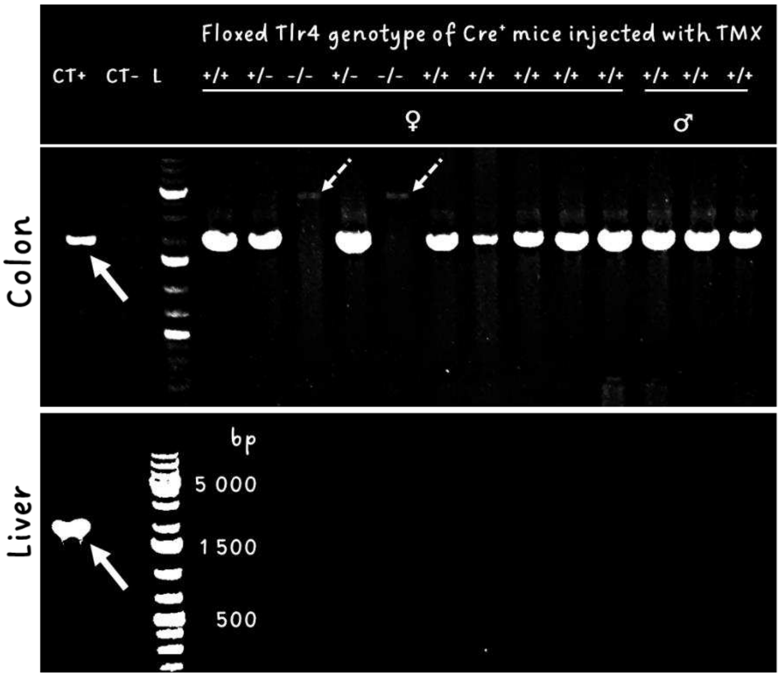


**Cre activity detection**

Villin specificity, Cre activity detection and impact of floxed genotype (homo or heterozygote) were evaluated by agarose 1% gel electrophoresis after PCR amplification.

Full arrows indicate excised site/Cre activity (1 768 bp) and slight dotted arrows indicated the full sequence (4 279 bp) of non floxed TLR4 sequence.

### (D) Specificity of villin conditional Cre expression

| Genotype | Cre activity (detection in PCR of excised site: + → detection corresponding to Cre activity; - → no detection; +/- → poor detection) | | | | | | | | | | |
|---|---|---|---|---|---|---|---|---|---|---|---|
| | Hypothalamus | Liver | Kidney | Intestins | | | | | | | |
| | | | | Proximal duodenum | Proximal jejunum | Middle of jejunum | Proximal ileum | Distal ileum | Proximal colon | Middle of colon | Distal colon |
| | | | | 1 | 2 | 3 | 4 | 5 | 6 | 7 | 8 |
| Cre + fl TLR4 +/+ | - | - | - | + | + | + | + | + | + | + | + |
| Cre + fl TLR4 +/- | - | - | - | + | + | + | + | + | + | + | + |
| Cre + fl TLR4 -/- | - | - | - | - | - | - | - | - | - | - | - |
| Cre + fl TLR4 +/- | - | - | - | + | + | + | + | + | + | + | + |
| Cre + fl TLR4 -/- | - | - | - | - | - | - | - | - | - | - | - |
| Cre + fl TLR4 +/+ | - | - | - | +/- | + | + | + | + | + | + | + |
| Cre + fl TLR4 +/+ | - | - | - | +/- | + | + | + | + | + | + | + |
| Cre + fl TLR4 +/+ | - | - | - | +/- | + | + | + | + | + | + | + |
| Cre + fl TLR4 +/+ | - | - | - | +/- | + | + | + | + | + | + | + |
| Cre + fl TLR4 +/+ | - | - | - | +/- | + | + | + | + | + | + | + |
| Cre + fl TLR4 +/+ | - | - | - | - | + | + | + | + | + | + | + |
| Cre + fl TLR4 +/+ | - | - | - | +/- | + | + | + | + | + | + | + |
| Cre + fl TLR4 +/+ | - | - | - | +/- | + | + | + | + | + | + | + |

**Table of Cre activity signal in different tissues and mice genotype, 17 days after 5 daily 1 mg tamoxifen injections.**

#### Figure S2

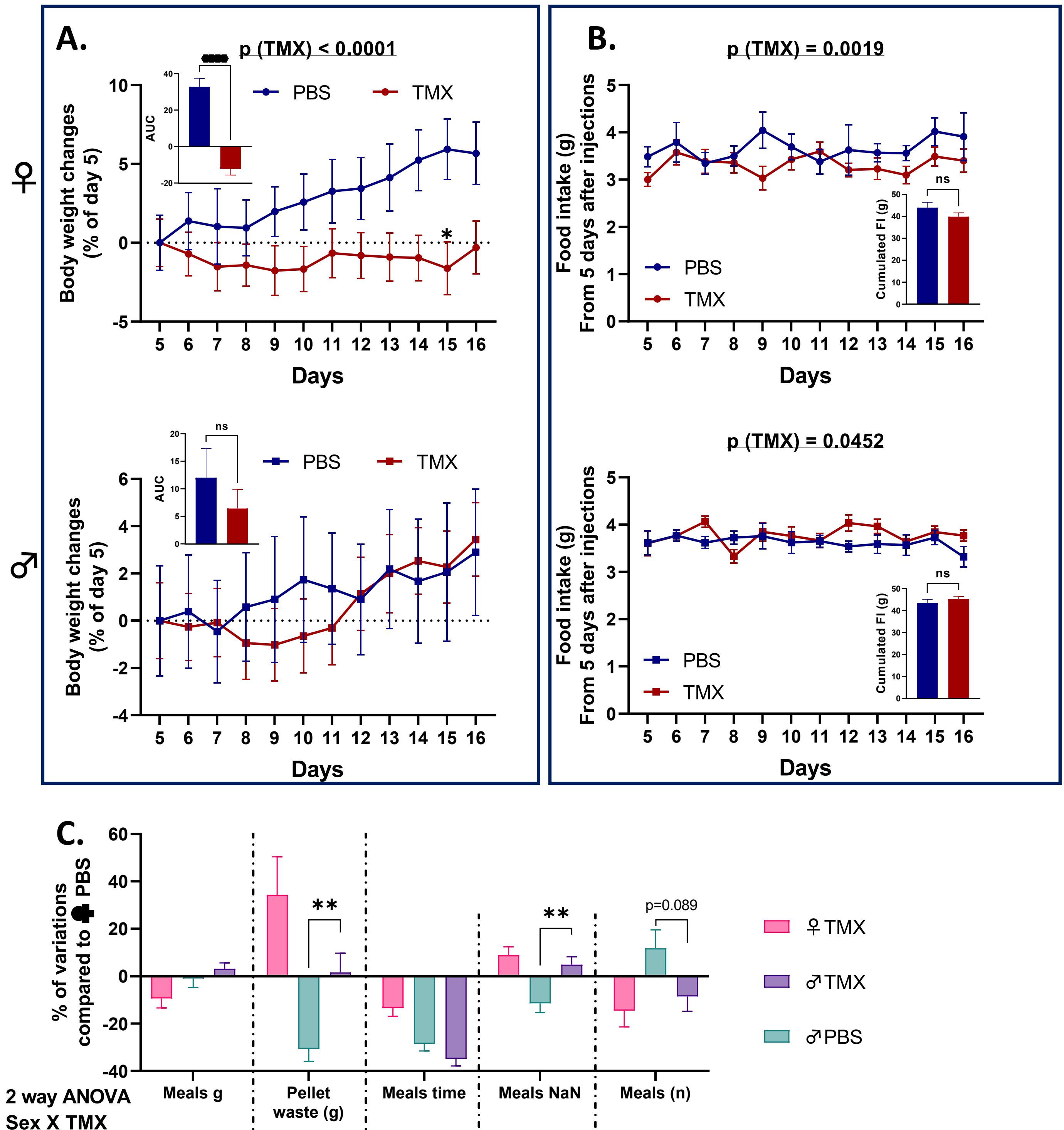


| 2 way ANOVA Sex X TMX | Meals g | Pellet waste (g) | Meals time | Meals NaN | Meals (n) |
|---|---|---|---|---|---|
| Int | | | | | |
| Sex | | * | * | | |
| TMX | | * | * | * | * |

#### Figure S3

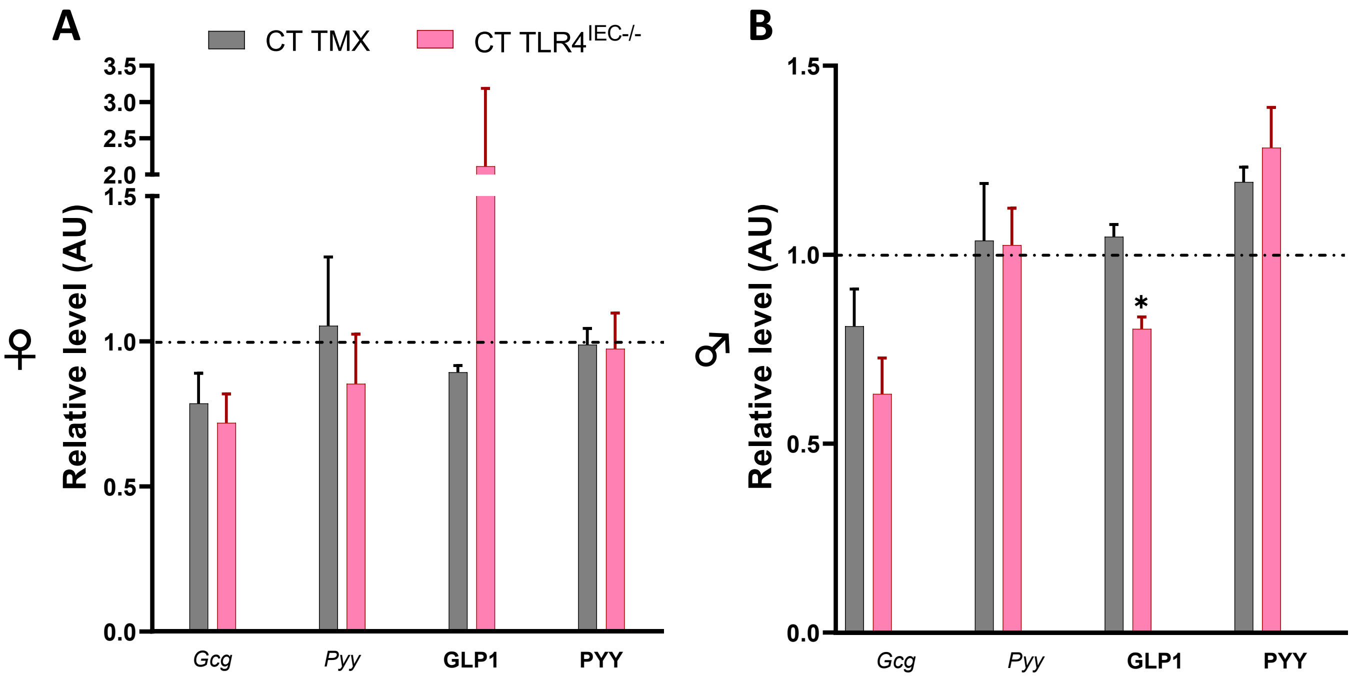
A
B
CT TMX
CT TLR4IEC-/-
Relative level (AU)
♀
♂
3.5
3.0
2.5
2.0
1.5
1.0
0.5
0.0
*
Gcg
Pyy
GLP1
PYY

**Figure S4**

**A**

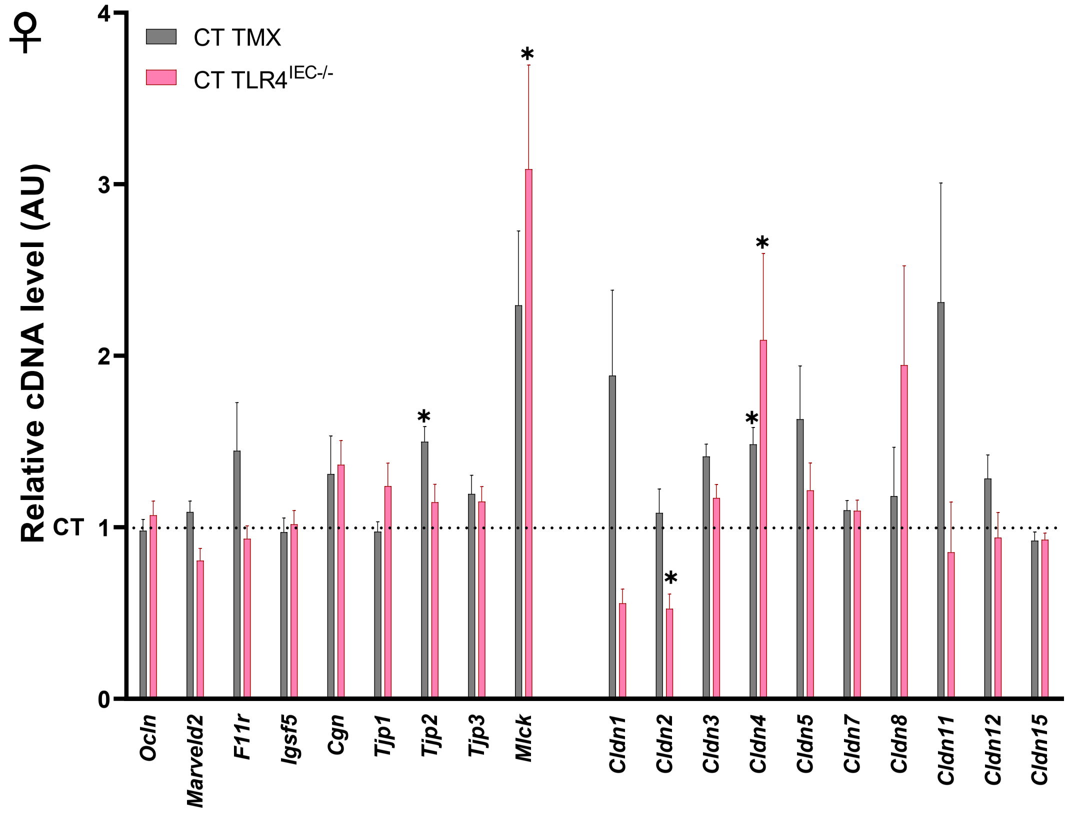
♀
Relative cDNA level (AU)
CT TMX
CT TLR4IEC-/-
CT
0
1
2
3
4
*
Ocln
Marveld2
F11r
Igsf5
Cgn
Tjp1
Tjp2
Tjp3
Mlck
Cldn1
Cldn2
Cldn3
Cldn4
Cldn5
Cldn7
Cldn8
Cldn11
Cldn12
Cldn15

**B**

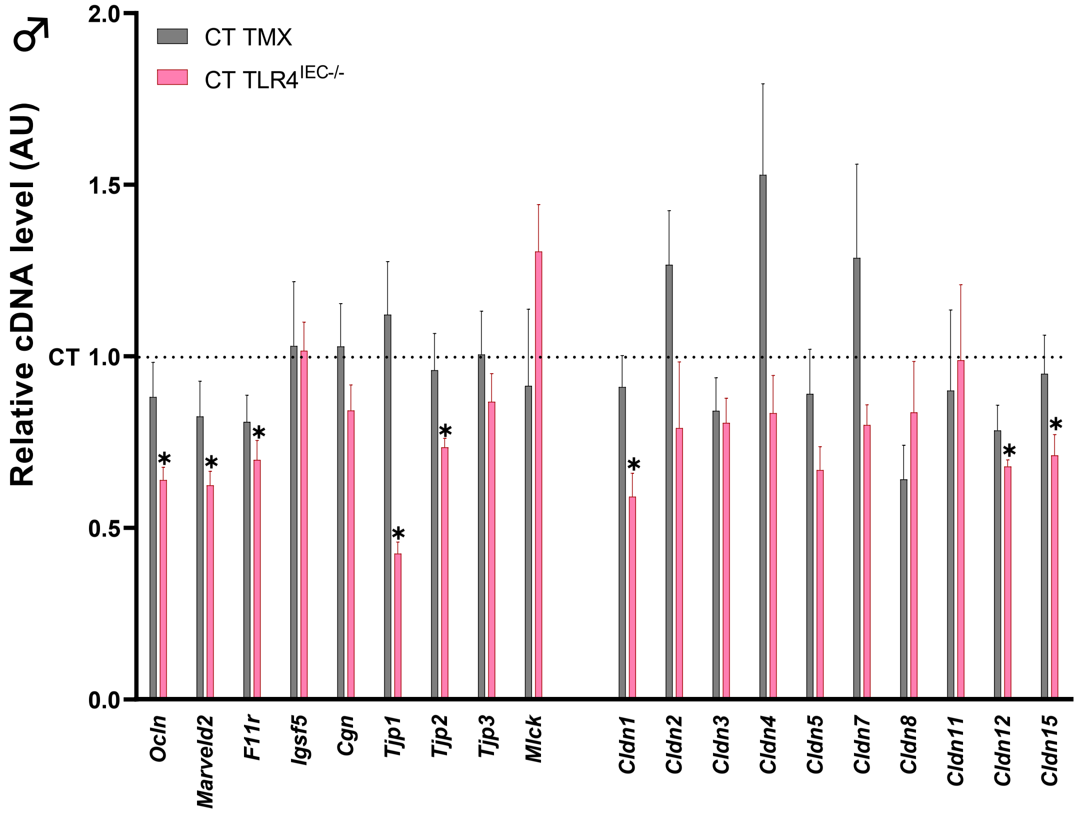
♂
Relative cDNA level (AU)
CT TMX
CT TLR4IEC-/-
CT
0.0
0.5
1.0
1.5
2.0
*
Ocln
Marveld2
F11r
Igsf5
Cgn
Tjp1
Tjp2
Tjp3
Mlck
Cldn1
Cldn2
Cldn3
Cldn4
Cldn5
Cldn7
Cldn8
Cldn11
Cldn12
Cldn15

Figure S5

A

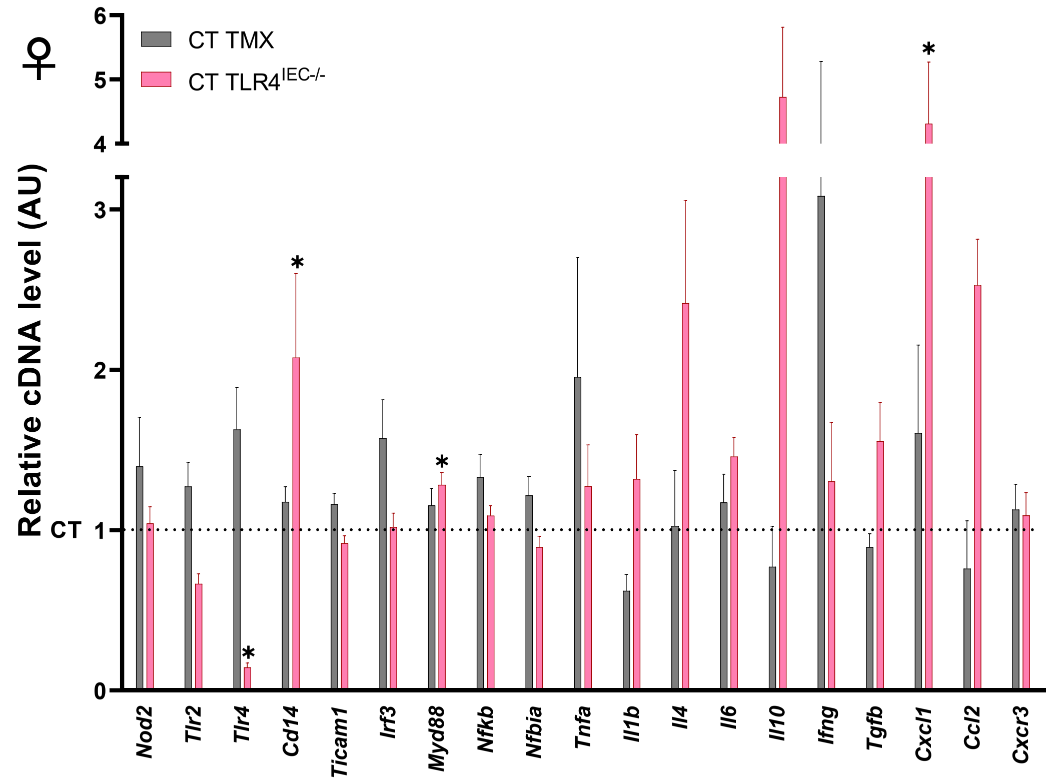
♀
CT TMX
CT TLR4IEC-/-
Relative cDNA level (AU)
CT
0
1
2
3
4
5
6
*
Nod2
Tlr2
Tlr4
Cd14
Ticam1
Irf3
Myd88
Nfkb
Nfbia
Tnfa
Il1b
Il4
Il6
Il10
Ifng
Tgfb
Cxcl1
Ccl2
Cxcr3

B

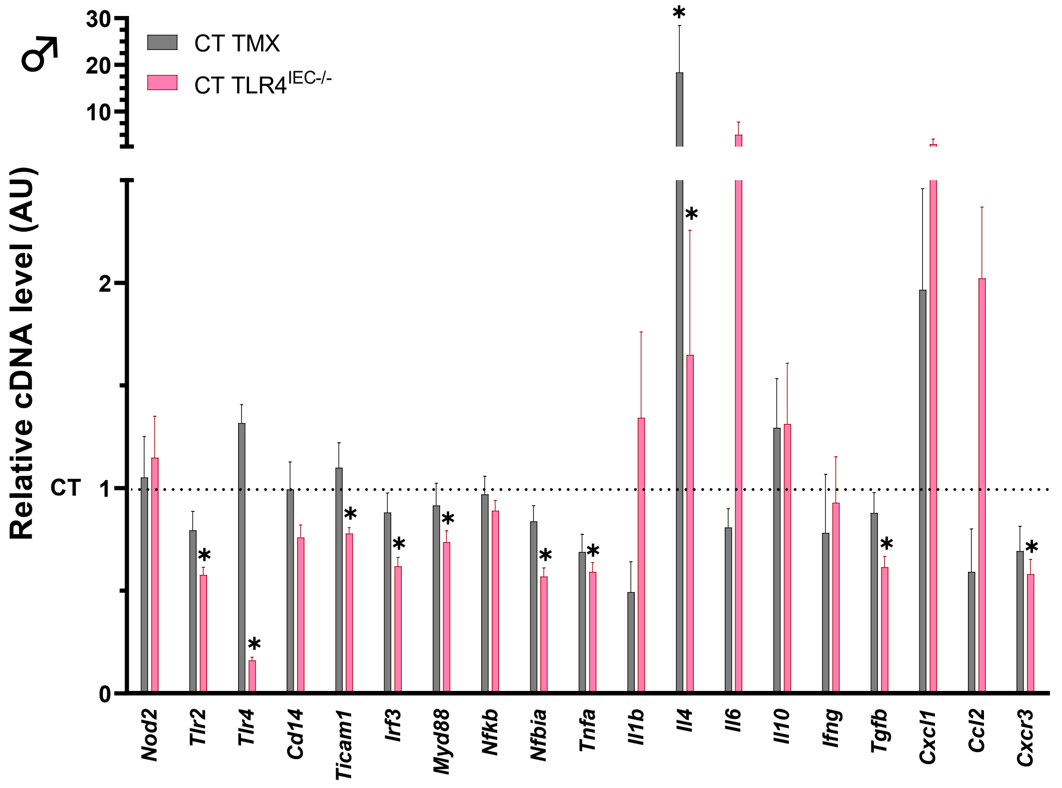
♂
CT TMX
CT TLR4IEC-/-
Relative cDNA level (AU)
CT
0
1
2
10
20
30
*
Nod2
Tlr2
Tlr4
Cd14
Ticam1
Irf3
Myd88
Nfkb
Nfbia
Tnfa
Il1b
Il4
Il6
Il10
Ifng
Tgfb
Cxcl1
Ccl2
Cxcr3

### Figure S6

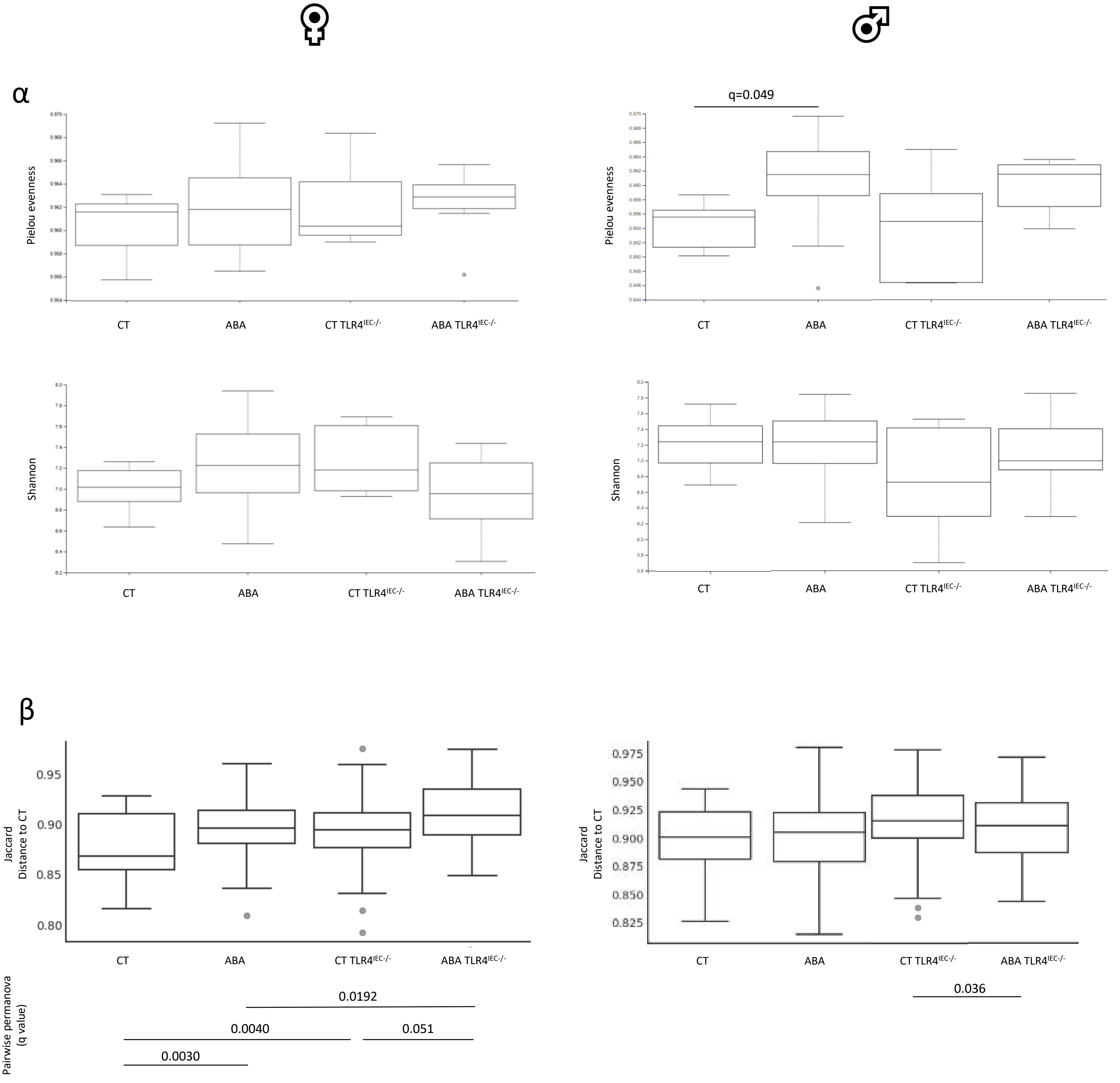
α
q=0.049
Pielou evenness
CT
ABA
CT TLR4$^{IEC-/-}$
ABA TLR4$^{IEC-/-}$
Shannon
β
Jaccard Distance to CT
Pairwise permanova (q value)
0.0192
0.0040
0.051
0.0030
0.036

#### Figure S7

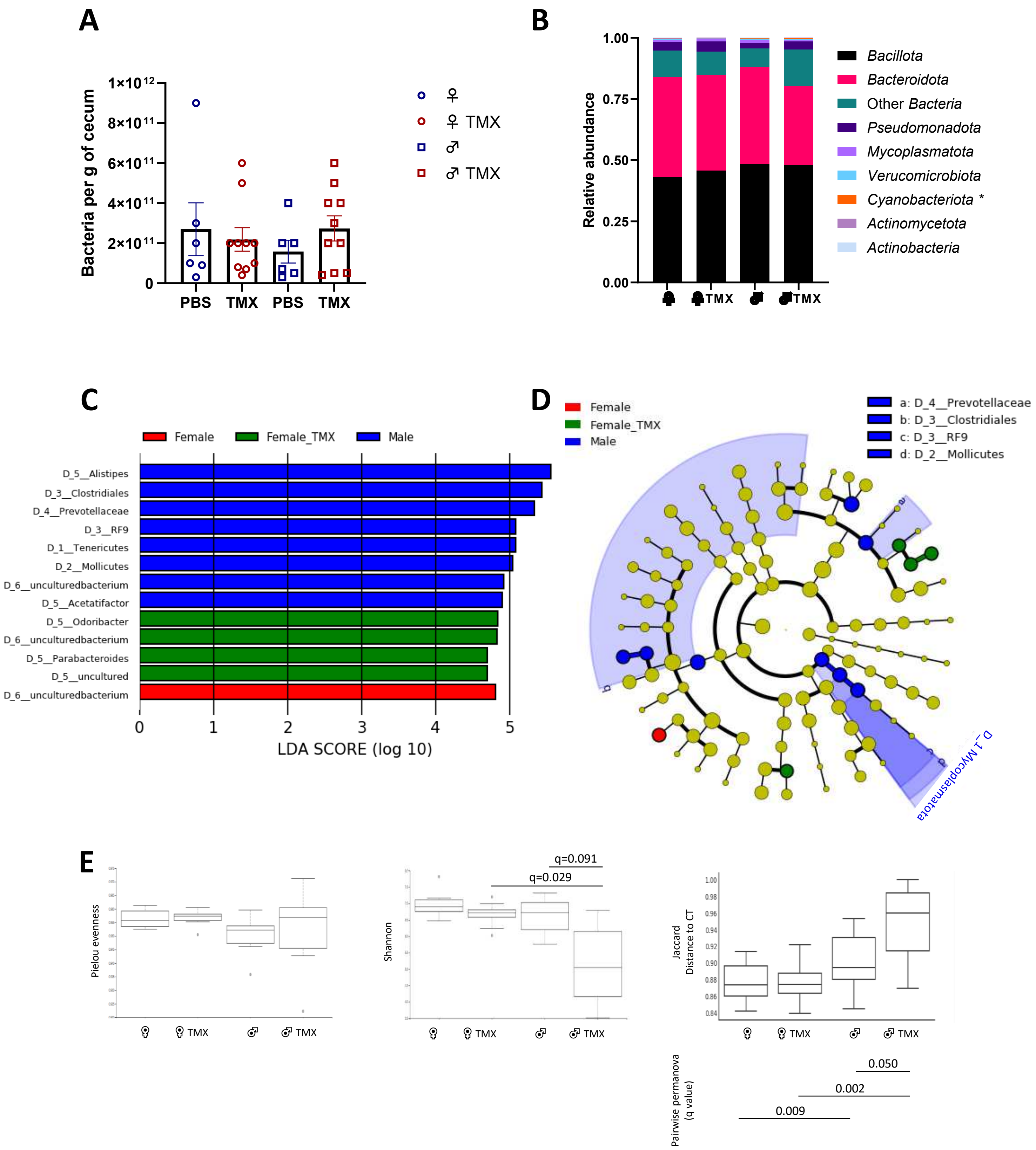

A
Bacteria per g of cecum
1×10^12
8×10^11
6×10^11
4×10^11
2×10^11
0
PBS
TMX
PBS
TMX
♀
♀ TMX
♂
♂ TMX
B
Relative abundance
1.00
0.75
0.50
0.25
0.00
Bacillota
Bacteroidota
Other Bacteria
Pseudomonadota
Mycoplasmatota
Verucomicrobiota
Cyanobacteriota *
Actinomycetota
Actinobacteria
C
Female
Female_TMX
Male
D_5__Alistipes
D_3__Clostridiales
D_4__Prevotellaceae
D_3__RF9
D_1__Tenericutes
D_2__Mollicutes
D_6__unculturedbacterium
D_5__Acetatifactor
D_5__Odoribacter
D_6__unculturedbacterium
D_5__Parabacteroides
D_5__uncultured
D_6__unculturedbacterium
LDA SCORE (log 10)
D
Female
Female_TMX
Male
a: D_4__Prevotellaceae
b: D_3__Clostridiales
c: D_3__RF9
d: D_2__Mollicutes
D_1_Mycoplasmatota
E
Pielou evenness
Shannon
q=0.091
q=0.029
Jaccard Distance to CT
1.00
0.98
0.96
0.94
0.92
0.90
0.88
0.86
0.84
TMX
Pairwise permanova (q value)
0.050
0.002
0.009
